# Mucin-mimetic glycan arrays integrating machine learning for analyzing receptor pattern recognition by influenza A viruses

**DOI:** 10.1101/2021.04.17.440161

**Authors:** Taryn M. Lucas, Chitrak Gupta, Meghan O. Altman, Emi Sanchez, Matthew R. Naticchia, Pascal Gagneux, Abhishek Singharoy, Kamil Godula

**Affiliations:** Department of Chemistry and Biochemistry, University of California San Diego, 9500 Gilman Drive, La Jolla, CA 92093; Department of Pathology, Division of Comparative Pathology and Medicine, University of California San Diego, 9500 Gilman Drive, La Jolla, CA 92093; Glycobiology Research and Training Center, University of California San Diego, 9500 Gilman Drive, La Jolla, CA 92093; School of Molecular Sciences, University, Tempe, AZ 85281; Biodesign Institute, Arizona State University, Tempe, AZ 85281

**Keywords:** Influenza A, hemagglutinin, mucin, glycan array, receptor pattern, machine learning

## Abstract

Influenza A viruses (IAVs) exploit host glycans in airway epithelial mucosa to gain entry and initiate infection. Rapid detection of changes in IAV specificity towards host glycan classes can provide early indication of virus transmissibility and infection potential. IAVs use hemagglutinins (HA) to bind sialic acids linked to larger glycan structures and a switch in HA specificity from α2,3-to α2,6-linked sialoglycans is considered a prerequisite for viral transmission from birds to humans. While the changes in HA structure associated with the evolution of binding phenotype have been mapped, the influence of glycan receptor presentation on IAV specificity remains obscured. Here, we describe a glycan array platform which uses synthetic mimetics of mucin glycoproteins to model how receptor presentation in the mucinous glycocalyx, including glycan type and valency of the glycoconjugates and their surface density, impact IAV binding. We found that H1N1 virus produced in embryonated chicken eggs, which recognizes both receptor types, exclusively engaged mucin-mimetics carrying α2,3-linked sialic acids in their soluble form. The virus was able utilize both receptor structures when the probes were immobilized on the array; however, increasing density in the mucin-mimetic brush diminished viral adhesion. Propagation in mammalian cells produced a change in receptor pattern recognition by the virus, without altering its HA affinity, toward improved binding of α2,6-sialylated mucin mimetics and reduced sensitivity to surface crowding of the probes. Application of a support vector machine (SVM) learning algorithm as part of the glycan array binding analysis efficiently characterized this shift in binding preference and may prove useful to study the evolution of viral responses to a new host.

## INTRODUCTION

The periodic emergence of new respiratory viruses capable of spreading across the human population continues to exact a significant toll on human life and the global economy. The novel coronavirus, SARS-Cov2, which is responsible for the ongoing global COVID-19 pandemic,^1^ provides a stark example of the risks of zoonotic virus adaptation to our society. Other animal pathogens, such as avian Influenza A viruses (IAVs), continuously pose a threat of crossing to human hosts and require close monitoring.^2^ Many respiratory viruses, including IAVs, utilize specific glycan receptors on airway epithelial cells to initiate entry and replication.^3^ Characterization of the glycan-binding phenotype of IAVs may provide an early indicator of increased infection potential.^4,5,6^

IAVs carry two types of glycoproteins in their viral coat with specificity for terminal sialic acid modifications on cell surface glycans – the receptor-binding hemagglutinins (HAs) and the receptor-destroying neuraminidases (NAs).^7,8,9^ The configuration of the sialic acid glycosidic bond linkage to the underlying glycans in glycoproteins and glycolipids plays a prominent role in defining IAV host specificity (**Fig 1A**). According to the prevailing paradigm,^4,10^ avian viruses preferentially recognize α2,3-linked sialic acids abundant in the gastrointestinal tract of birds, while human IAVs have affinity for α2,6-sialosides expressed on lung epithelial cells in our upper airways. A switch in HA specificity from α2,3-to α2,6-linked sialic acids is associated with increased infection and transmission in humans.^4,11,12^

**Figure 1.**
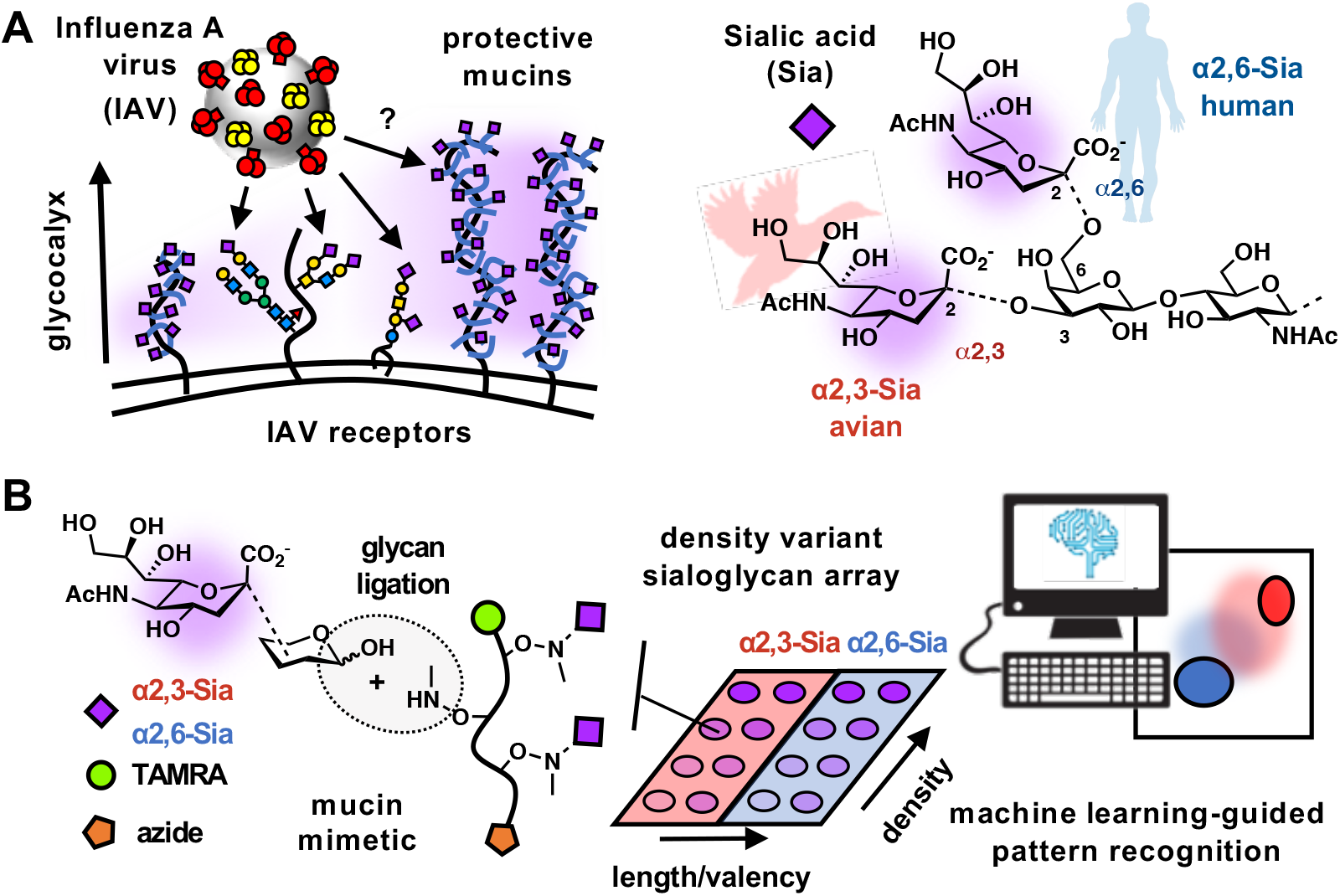
Machine learning-enabled glycomimetic array platform for assessing receptor pattern recognition by Influenza A viruses (IAVs). A) IAVs begin their infection cycle by binding to sialylated host glycans, but these receptors are also present on mucins which have a proposed protective function. Avian and human IAVs show distinct preferences for the binding of α2,3- and α2,6-sialoglycan receptors. B) Glycopolymers, which mimic the architecture and composition of mucins, were used to build models of the mucinous glycocalyx on microarrays. A supervised vector machine (SVM) learning algorithm enabled analysis of viral binding response to changing receptor patterns in the synthetic glycocalyx arrays.

Glycomics screens^13,21^ and cell-based studies using glycosylation mutants^14,15^ have revealed that, in addition to a particular sialic acid linkage configuration, IAVs can also discriminate between distinct glycan classes and glycoconjugate types (i.e., *N*- and *O*-glycosylated proteins and glycolipids). Spatial combinations of these sialylated glycoconjugates give rise to three-dimensional, hierarchically organized receptor patterns in the host cell glycocalyx that determine the specificity and avidity of IAV binding. Glycan arrays, which present ensembles of chemically defined glycans printed and immobilized on glass substrates, are routinely used to analyze viral HA-receptor specificity.^16,17,18,19,20^ However, a recent cross-comparison between glycan composition of *ex vivo* human lung and bronchus tissues with glycan array binding data pointed to a limited ability of the arrays to predict infection events.^21^ This indicates that the current platforms may not accurately recapitulate the receptor presentation in the glycocalyx environment as encountered by viruses at the mucosal epithelium.

The mucosal epithelial cell glycocalyx is dominated by membrane-tethered mucins (MUCs), which are large, heavily glycosylated proteins projecting tens to hundreds of nanometers above the cell surface (**Fig 1A**).^22,23^ Mucins carry primarily, but not exclusively, *O*-glycans linked to tracks of serine and threonine residues within the core protein. As much as 80% of mucin mass derives from glycans, giving these glycoproteins an extended semi-flexible bottlebrush form.^24^ The *O*-glycans in mucins are frequently terminated with sialic acids that can serve as IAV receptors;^25^ however, epithelial mucins are believed to primarily provide protection against infection. Mucins can serve as decoys, which shed from the cell surface upon virus binding,^22^ or assemble into dense extended glycoprotein brushes that restrict virus access to apical membrane receptors and interfere with internalization.^26,27^ Due to the extension of these proteins within the glycocalyx and their propensity for carrying sialic acid, mucins are most likely the first, and likely non-productive, site of virus attachment in its infection cycle.^28^ Interestingly, the IAV subtype, H1N1, was found to colocalize with some (i.e., MUC1) but not other (i.e., MUC13 and MUC16) mucins on the surfaces of A549 lung epithelial cells,^29^ revealing a preference of the virus for distinct mucin family members within the same cell produced by a shared glycosylation machinery. The type of mucin and its presentation at the cell surface is likely to influence initial viral interactions at the epithelium and determine the course of infection. A more complete understanding of IAV interactions at the mucosal glycocalyx may, thus, provide a more accurate assessment of the potential of IAVs to infect human hosts.

Here, we report the development of an array platform, in which synthetic mucin-mimetics are used to model the mucosal epithelial cell glycocalyx, to evaluate receptor pattern recognition by IAVs (**Fig 1B**). We developed supervised machine learning to identify and analyze effects of variations in glycan receptor type, mucin mimetic valency, nanoscale dimensions, and crowding in the glycocalyx models on shifts in the binding specificity of IAVs. We found that mucin-like polyvalent presentations of α2,3- and α2,6-sialoglycans differentially impacted their recognition by the H1N1 virus and that surface crowding of the glycoconjugates limited viral adhesion, consistent with the proposed protective functions of mucins in the airway epithelium. The mucin mimetic arrays also revealed an evolution of receptor pattern recognition by IAVs produced in both, avian and mammalian cells, which could be characterized through machine learning.

## RESULTS AND DISCUSSION

### Construction of glycopolymers for mucin-like glycan receptor presentation

To model the mucinous glycocalyx environment in glycan arrays, we have devised a method for generating synthetic glycopolymers (**GPs**) that replicate key structural features of mucins (i.e., polyvalent glycans displayed along extended linear polypeptide chains) while allowing for tuning of the mimetic size and glycosylation pattern (**Fig 2A**). Using the reversible addition-fragmentation chain transfer (RAFT) polymerization, we have generated a collection of mucin mimetics of increasing length glycosylated with α2,3- and α2,6-siallyllactose trisaccharides (**α2**,**3-SiaLac** and **α2**,**6-SiaLac**) as model avian and human IAV receptors, respectively. The polymers were terminated with an azide functionality and used either as soluble probes or covalently grafted on cyclooctyne-coated glass via the strain-promoted alkyne-azide cycloaddition (SPAAC) reaction to a mucin-like glycocalyx display.^30^ A tetramethylrhodamine (TAMRA) fluorophore was appended to the opposing chain end to allow for characterization of mucin mimetic density on the arrays.

**Figure 2.**
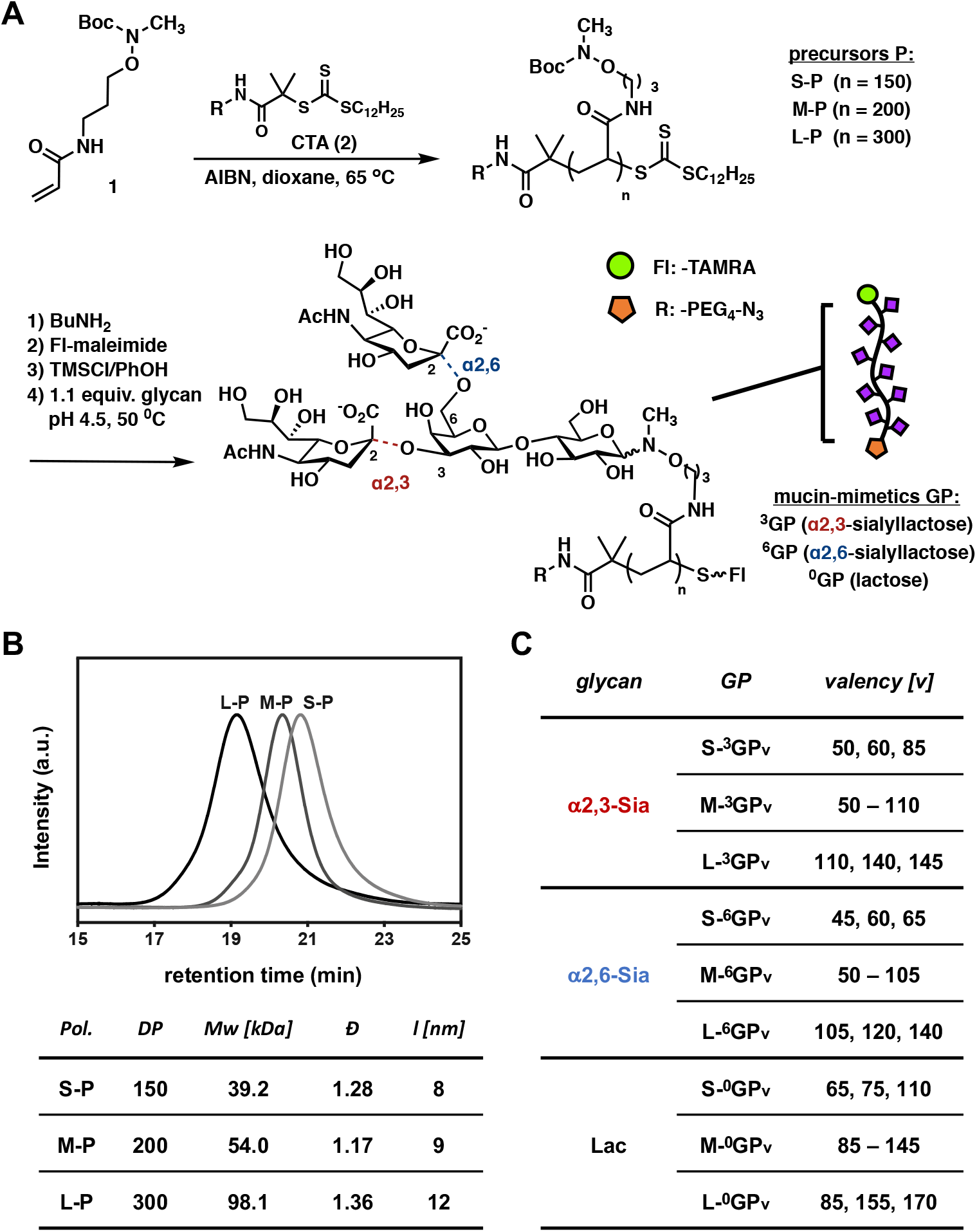
Generation of mucin mimetic probes. A) Fluorescently-labeled azide-terminated short (**S**), medium (**M**) and long (**L**) mucin-mimetic glycopolymers **GP** were generated via RAFT polymerization. B) Size exclusion chromatography (SEC) analysis of the polymeric precursors **P**. C) The naming scheme for the **GPs** indicates the polymer backbone length (**S**-, **M**-, and **L**-), sialic acid linkage type (superscripts 3 and 6, and Ø designate **α2**,**3-SiaLac, α 2**,**6-SiaLac**, and **Lac**, respectively), and glycan valency (final subscript).

The mucin mimetic synthesis began with the polymerization of a Boc-protected *N*-methylaminooxypropyl acrylamide monomer (**1**) in the presence of a chain transfer agent (CTA, **2**) and a radical initiator (AIBN) to generate a set of azide-terminated short (**S**, DP ~ 150), medium (**M**, DP ~ 200) and long (**L**, DP ~ 300) polymeric precursors, **P** (**Fig 2A**). Size exclusion chromatography analysis (SEC, **Fig 2B**) confirmed good control over the target molecular weight (Mw) and dispersity (Đ) of the polymers. Next, the trithiocarbonate end groups in polymers **P** were removed by aminolysis and the newly exposed thiol groups were capped with TAMRA-maleimide (**Fig 2A** and **Scheme S1**). The fluorophore labeling efficiency was determined for each polymer by UV-VIS spectrometry and ranged between 6-30% (**Tables S3**). Side-chain Boc-group deprotection in the presence of phenol and trimethylsilyl chloride (TMSCl)^31^ followed by conjugation of the released *N*-methylaminooxy groups with reducing glycans under acidic conditions completed the synthesis of the mucin mimetic glycopolymers **GP** (**Fig 2A**)

The mucin mimetic library was comprised of 27 short (**S**), medium (**M**) and long (**L**) sialylated glycopolymers, ^**3**^**GP** and ^**6**^**GP**, decorated with increasing amounts of the trisaccharides, α2,3-**SiaLac** and α2,6-**SiaLac**, respectively (**Fig 2C** and **Table S3**). In addition, we generated 11 control polymers lacking sialic acid modifications (^**Ø**^**GP**) displaying only the lactose disaccharide (**Lac, Fig 2C** and **Table S3**). The extent of glycosylation for all polymers was determined by ^1^H NMR spectroscopy (**SI Appendix 1**) and varied according to glycan structure. Treatment with 1.1 equiv. of glycan per polymer sidechain was sufficient to achieve maximum polymer glycosylation of ~ 70% for **Lac** and ~ 45% for the negatively charged **α2**,**3-SiaLac** and **α2**,**6-SiaLac** (**Table S3**). The use of sub-stoichiometric amounts of glycans enabled tuning of glycan valency in the mucin mimetics (**Fig 2C** and **Fig S1**). The mucin mimetic lengths (*l*) were estimated to range from ~ 8 nm to 12 nm according to their DP using a method by Miura *et al*. for calculating theoretical end-to-end distances in sialylated glycopolymers (**Fig 2B** and **Equation S1**).^32^

### The oligomeric plant lectins, WGA and SNA, show distinct binding behavior in mucin-like receptor displays

In the airways, cell surface-associated mucins are organized into a dense, brush-like glycocalyx, which projects tens to hundreds of nanometers above the epithelial cell surface.^22^ To gain insights into glycan receptor recognition by proteins and pathogens at the mucosal interface, we modeled the mucinous glycocalyx in arrays by printing mucin mimetic glycopolymers **GP** on cyclooctyne-functionalized glass (**Fig 3A**). In addition to varying the structure and glycosylation of the glycopolymer probes, we also modulated their surface crowding by increasing their concentration (c_***GP***_) from 1 to 10 µM in the printing buffer (PBS supplemented with 0.05% Tween-20, pH = 7.4). The fluorescent TAMRA labels introduced synthetically into the probes were used to establish the surface grafting efficiency (**Fig S2A and C**) and the glycan receptor density (**Fig S2C**) for each polymer condition. The printing conditions yielded spots of uniform morphology (**Fig 3A**) with linear increase in polymer density across the employed concentrations regardless of polymer size or glycosylation (**Fig S3** and **Fig S4**).

**Figure 3.**
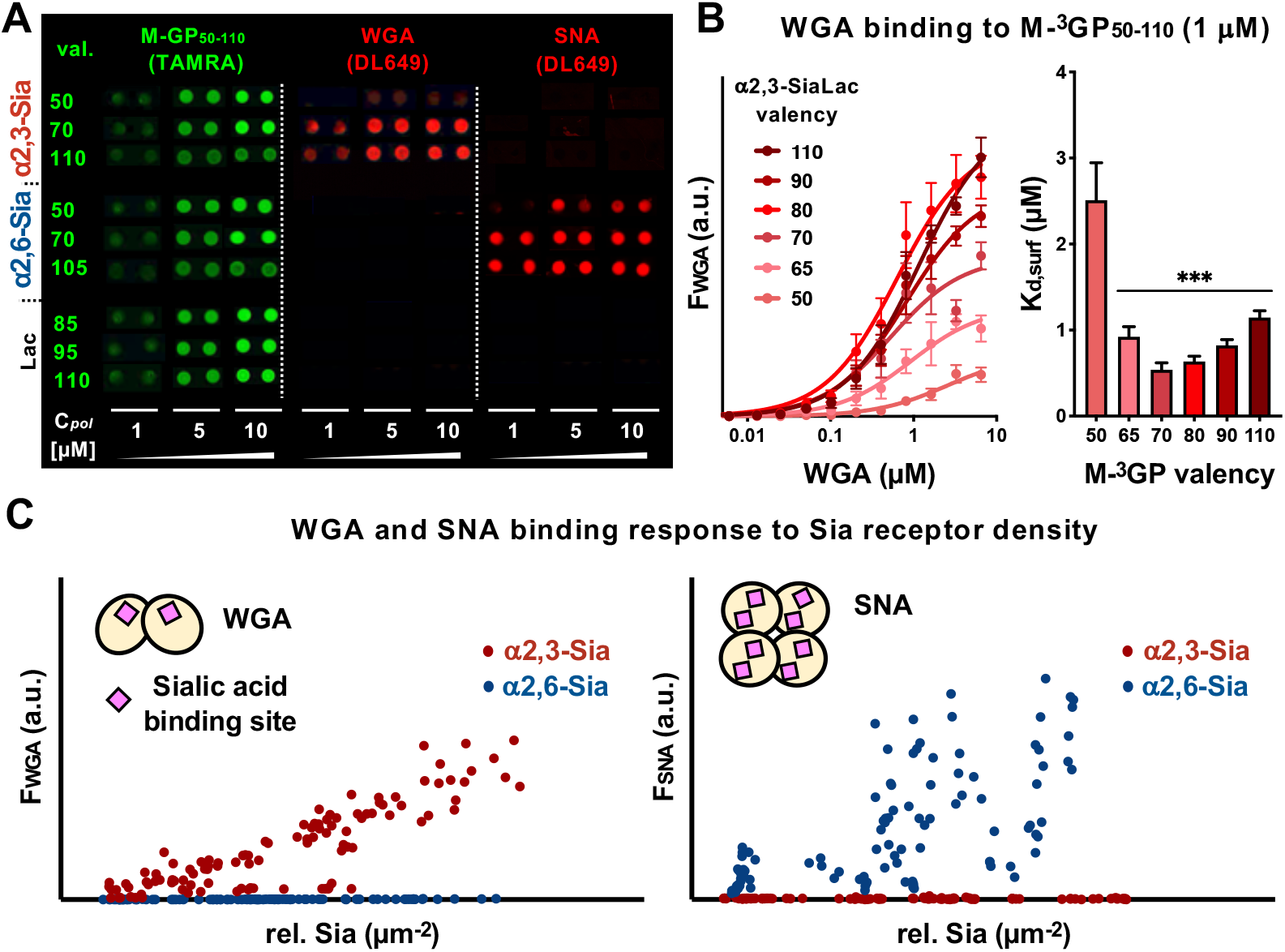
Construction and validation of mucin mimetic arrays. A) Representative composite images of density-variant arrays of fluorescent mucin-mimetic glycopolymers **GP** (TAMRA, green) probed with Daylight649-labeled SNA and WGA lectins. Each condition is represented as a duplicate. Full array scans are provided in **Fig S5**. B) Binding isotherms and associated apparent surface dissociation constants (*K*_*D,surf*_) for binding of WGA to medium-sized mucin mimetics **M-** ^**3**^**GP**_**50-110**_ with increasing **α2**,**3-SiaLac** valency printed at low surface density (c_***GP***_ = 1 μM, ***p <0.0005 or greater). C) Binding responses of WGA and SNA to increasing glycan receptor density on the array. The dimeric WGA lectin binding is directly proportional to glycan density, whereas the tetrameric SNA lectin exhibits a more complex binding pattern. Insets represent graphical representation of lectin oligomeric state and orientation of sialic acid binding sites based on crystallographic data analysis (**Figs S8** and **S9**).

To confirm selective recognition of the mucin mimetics based on to the structure of their pendant glycans, the arrays were probed with Dylight649-labeled lectins wheat germ agglutin (WGA) and *Sambucus nigra* agglutinin (SNA) with known preference for α2,3- and α2,6-linked sialic acids, respectively (**Fig 3A** and **Fig S5**).^33,34^ To obtain quantitative assessment of lectin binding to the mucin mimetics, we probed the arrays with increasing concentrations of the lectins to establish binding isotherms and extract apparent surface dissociation constants (*K*_*D,surf*_) (**Fig 3B** and **Fig S6**). WGA binding to the medium sized **α2**,**3-SiaLac** polymers, **M**-^**3**^**GP**_**50-110**,_ printed at low surface density (c_***GP***_ = 1µM) showed valency-dependent binding with autoinhibition at the highest valencies caused by glycan crowding on the polymer backbone. This behavior is frequently observed for lectin binding to glycopolymer probes in solution.^35^ The low polymer printing concentration produced probe spacing on the array surface that allowed for the measurement of lectin binding responses to the underlying glycoconjugate architecture. Increasing the concentration of the polymers resulted in denser mucin mimetic arrays, attenuated WGA responsiveness to **α2**,**3-SiaLac** valency of the individual probes, and increased overall avidity of the dimeric lectin toward the receptor display (**Fig S6** and **Table S4**). Our attempts to establish similar binding profiles for SNA were not successful due to protein aggregation at concentrations needed to reach saturation binding (**Fig S7**).

Collecting thermodynamic binding data for each lectin-probe combination in the array can be time consuming and may not be possible for some lectins, as was the case for SNA. Simplified plots of lectin binding in response to changing relative sialoglycan density in the arrayed glycopolymer spots provide a convenient way to discern different binding modes of the proteins. Using this analysis, we observed that the binding response of WGA to changing glycan density was generally linear while SNA showed a less correlated binding pattern indicative of contributions from higher-order binding interactions, such crosslinking of neighboring glycopolymers on the array (**Fig 3C**). Analysis of crystallographic data for WGA and SNA provide a structural basis for their differences in crosslinking capacity (**Fig S8** and **Fig S9**). WGA exists as a dimer with two sialic acid binding domains separated by 3.9 nm and positioned on the same face of the protein.^34^ This arrangement reasonably favors WGA binding to glycans presented on the same mucin mimetic and may be responsible for the largely linear relationship between receptor density and lectin binding response. By contrast, SNA can exist as either a monomer, dimer, or tetramer.^36^ Each monomer contains two glycan binding sites that are directed outward on opposite the edges of the protein.^37^ The various oligomeric states and the orientation of the binding sites make SNA more likely to engage and crosslink multiple glycoconjugates on the surface producing the more complex binding behavior observed on the mucin mimetic arrays.

### H1N1 PR8 virus shows linkage-specific differences in binding to mucin-like sialoglycan presentations

Pathogens, which utilize oligomeric lectins and adhesins for binding to cell-surface glycan receptors, may be sensitive to the presentation of glycan receptors at the mucosal barrier.^38^ We examined the binding of the H1N1 (A/Puerto Rico/8/1934 or PR8) virus to different presentations of sialoglycan receptors in our mucin mimetic platform (**Fig 4A**). The PR8 strain is a well-characterized, laboratory-adapted human IAV strain, which has the ability to recognize both avian and human sialic acid receptor structures.^39,40^ As such, it provides a useful model for assessing how receptor presentation may affect viral binding and selectivity.

**Figure 4.**
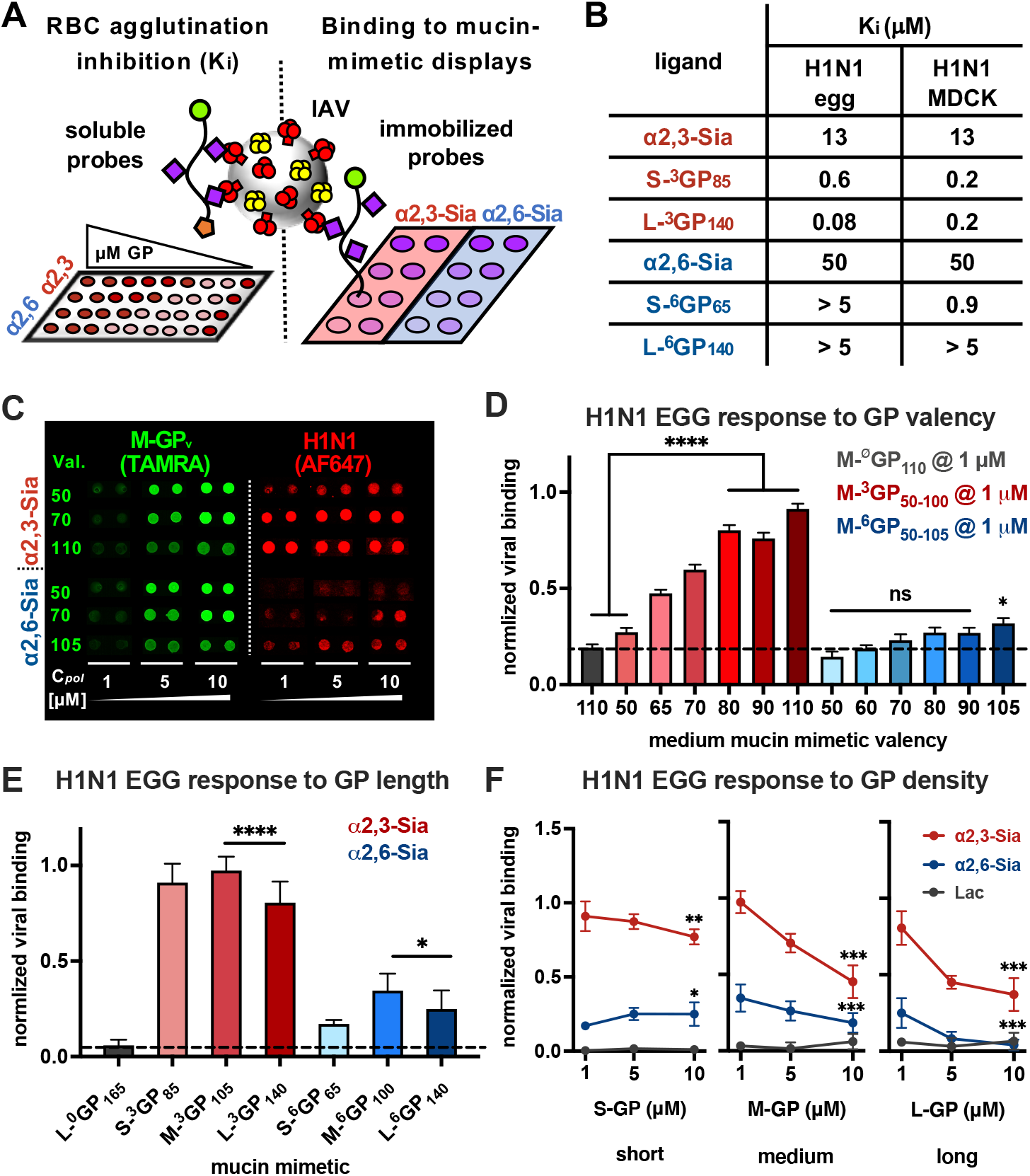
H1N1 EGG binding to mucin-mimetic displays of sialoglycan receptors. A) Red blood cell agglutination assays and array screens were used to probe the interactions of H1N1 produced in embryonated chicken eggs (H1N1 EGG) with soluble and surface bound mucin mimetics. B) Inhibitory activity, K_*i*_, of soluble glycan receptors **α2**,**3-SiaLac** and **α2**,**6-SiaLac** and mucin mimetic glycopolymers **GP** in RBC agglutination assays expressed as the minimal ligand concentration needed to prevent cell aggregation. C) Representative composite images and bar graph representation of H1N1 EGG virus (red) binding to medium-sized mucin mimetics **M-GP** (green) printed at low surface density (c_GP_ = 1 µM) according to glycan receptor valency. Each array condition is represented as a duplicate and full array images are included in **Figures S11, S12**, and **S14**. Values and error bars represent averages and standard deviations of experiments from 9 different arrays. Significance is based on viral binding to the **Lac** polymer control **M-** ^**Ø**^**GP**_**110**_ (black dashed line). E) H1N1 EGG binding to mucin mimetics of increasing length printed at low surface density (c_GP_ = 1 µM). Values and error bars represent averages and standard deviations of experiments from 6 different arrays. Significance was determined against **Lac** polymer control **L-**^**Ø**^**GP**_**165**_ (black dashed line). F) H1N1 EGG binding response to increasing crowding of mucin mimetics of all three lengths on the array surface. Values and error bars represent averages and standard deviations of experiments from 6 different arrays. Significance was determined against H1N1 binding at c_***GP***_ = 1 µM. (*p<0.05, **p<0.005, and ***p<0.0001)

H1N1, which was propagated in embryonated chicken eggs and henceforth labeled as H1N1 EGG, bound both receptor types in their soluble monovalent form, with ~ 4-fold preference for the **α2**,**3-SiaLac** isomer, as determined in red blood cell (RBC) agglutination inhibition assays (*K*_*i*,α**2**,**3**_ = 13 µM vs *K*_*i*,α**2**,**6**_ = 50 µM, **Fig 4B**). The array binding data mirrored this preference, while providing additional insights into the effects of receptor presentation on viral interactions (**Fig 4C-F**). H1N1 EGG virus binding to the medium size **α2**,**3-SiaLac** mucin mimetics **M-**^**3**^**GP**_**50-110**_ immobilized at low surface densities (c_***GP =***_ 1 µM) indicated enhanced viral capture with increasing receptor valency, with a valency threshold for binding above 50 **α2**,**3-SiaLac** residues and a plateau at ~ 80 glycans per polymer (**Fig 4D** and **Fig S11**). Shortening the polymer length while maintaining a high receptor valency above 80 (**S-**^**3**^**GP**_**85**_) had no negative effect on viral capture (**Fig 4E** and **Fig S12**). We observed some decrease in binding to the longest mucin mimetic **L-** ^**3**^**GP**_**140**_ compared to **M-**^**3**^**GP**_**105**_ despite its higher valency, presumably due to its increased chain conformational flexibility. RBC hemagglutination inhibition assays with soluble **α2**,**3-SiaLac** polymers ^**3**^**GP** confirmed the observed valency-dependent binding trend for H1N1 EGG (*K*_*i*,,**3GP**_ = 313 nM – 1.25 µM, **Fig S10**) and were consistent with prior reports using similar multivalent glycopolymers.^32^ In contrast to the behavior of the arrayed mucin mimetics, increasing the polymer length resulted in a more effective inhibition of RBC agglutination by H1N1 EGG in solution (*K*_*i*,**S-3GP**_ = 625 nM vs *K*_*i*, **L-3GP**_ = 78 nM, **Fig S10**). It should be noted that the increase in inhibitory capacity of the glycopolymers compared to the monovalent receptor can be accounted for based on glycan valency and concentration alone rather than avidity enhancements due to multivalency. In the case of the short polymer **S-**^**3**^**GP**_**85**_, when the total amount of glycan on the polymer is taken into account, the per glycan inhibitory activity (K_*i*,**S-3GP**_ x **α2**,**3-SiaLac** valency = 53 µM) was reduced compared to the free **α2**,**3-SiaLac** (K_*i*,α2,3-SiaLac_ = 13 µM). Lactose glycopolymers, ^Ø^**GP**, lacking sialic acids served as negative controls in both assays (**Fig 4** and **Fig S10**).

Glycopolymers carrying the **α2**,**6-SiaLac** receptor (**M-**^**6**^**GP**_**50-105**_) showed only a limited ability to engage H1N1 and required glycan valency above 90 to reach binding above background (**Fig 4D** and **Fig S10**). Extending the length of the mucin mimetic, again, resulted in a decrease in viral capture (**Fig 4E** and **Fig S12**). All of the **α2**,**6-SiaLac** polymers failed to inhibiting RBC hemagglutination by the virus over the range of tested polymer concentrations (*K*_*i*,**6GP >**_ 5 µM or 325 µM with respect to **α2**,**6-SiaLac, Fig S10**). Considering that monovalent **α2**,**6-SiaLac** can prevent RBCs agglutination (*K*_*i*,α**2**,**6** =_ 50 µM, **Fig 4B**), it appears that binding of H1N1 EGG to this glycan receptor is disfavored in the polyvalent glycopolymer presentation.

High levels of mucin expression on the surfaces of epithelial cells produces a dense glycoprotein brush, which has been proposed to restrict IAV access to membrane receptors necessary for infection.^26,27^ To examine the effects of polymer size and density on viral adhesion, we modelled glycocalyx crowding in our arrays by increasing the printing concentration of the mucin mimetics. We assayed H1N1 EGG binding to maximally glycosylated mucin mimetics of all three lengths arrayed at concentrations of 1, 5, and 10 µM (**Fig 4F** and **Fig S12**). The virus retained its overall preference for the **α2**,**3-SiaLac** probes across all surface densities; however, increased crowding of the polymers led to attenuated viral adhesion, which became more pronounced with increasing mucin mimetic length. Crowding of the **α2**,**6-SiaLac** glycopolymers both enhanced (**S-**^**6**^**GP**_**65**_) and inhibited (**L-**^**6**^**GP**_**140**_) viral adhesion depending on polymer length (**Fig 4F**). Our data show that, while the H1N1 EGG virus can utilize the less preferred **α2**,**6-SiaLac** receptors when presented in surface displays on short mucin mimetic scaffolds, increasing the length and density of the conjugates generally negatively impacted viral adhesion regardless of receptor type. Thus, crowding of mucins in the glycocalyx may not only shield underlying glycan receptors from the virus,^26^ but also limit viral adhesion to the heavily sialylated mucins themselves.

The observed differential H1N1 EGG binding to the mucin-like receptor displays according to sialic acid linkage type supports the distinct functions of secreted and membrane bound mucins comprising the airway mucosal barrier.^26,41^ Therein, secreted mucins produced by goblet cells and presenting primarily α2,3-sialic acid modifications serve as decoy receptors for viral capture and clearance. By contrast, the membrane-tethered mucins produced by epithelial cells display α2,6-linked sialic acid receptors and are thought to limit viral adhesion. The binding of H1N1 EGG to polyvalent **α2**,**3-SiaLac** mucin mimetics but not the **α2**,**6-SiaLac** analogs and the sensitivity of the virus to surface crowding of mucin mimetics carrying both receptor types would provide a rationale for the synergistic but mechanistically distinct functions of secreted and surface-bound mucins in limiting viral infection.

### H1N1 PR8 propagation in mammalian cells enhances interactions with α2,6-sialoglycans in mucin-like displays

Having established the influence of receptor organization in mucin glycocalyx-like displays on H1N1 adhesion, we set to analyze how the glycan binding phenotype of the virus may change depending on the host in which it is propagated. Such information may enhance existing viral surveillance by either eliminating glycan binding phenotype artifacts, which may be introduced during propagation of field-isolated viruses in the laboratory, or by establishing specific binding phenotype features associated with enhanced human transmission.

We performed a comparative binding analysis of H1N1 produced in Madin-Darby Canine Kidney (MDCK) cell culture in our mucin mimetic arrays. MDCK cells are a commonly used mammalian system for the propagation of IAVs in the laboratory. While the MDCK cell-derived H1N1 (H1N1 MDCK) binding to the soluble monovalent **α2**,**3-SiaLac** and **α2**,**6-SiaLac** glycans measured via RBC hemagglutination inhibition remained unchanged with ~ 4 fold preference for the α2,3-linked isomer, the virus exhibited improved ability to engage mucin mimetics carrying the **α2**,**6-SiaLac** receptors both in their soluble form (*K*_*i*,**S-6GP65**_ = 938 nM, **Fig 4B**) and on arrays (**Fig S13** and **Fig S14**). The increased ability to recognize α2,6-sialoglycans is expected for H1N1 propagated in MDCK cells. While the cells in the allantoic fluid of embryonated chicken eggs used for viral propagation display mainly α2,3-linked sialic acids, the surfaces of MDCK cells are populated by receptors with both linkages.^42^ During culture in MDCK cells, viruses can undergo selection for both receptor types, giving rise to virus population with an altered sialoglycan binding phenotype.^11,42^ However, our hemagglutination inhibition assays indicate that the altered glycan-binding phenotype of H1N1 MDCK does not arise from changes in hemagglutinin affinity for the individual monovalent receptors (**Fig 4B**).

### Machine learning enables analysis of receptor pattern recognition by H1N1 viruses

Next, we set to more systematically analyze the viral binding phenotypes of H1N1 for features that are conserved between viruses produced in embryonated eggs and those propagated in MDCK cells. While the focused analysis of some of the key features of glycan presentation in the mucin mimetic array as described above (e.g. polymer size, valency, or surface density) was informative, the multidimensionality of receptor presentation on the array makes comprehensive analysis challenging and time consuming. Correlating binding responses of SNA and WGA lectins with glycan density in the array, regardless of polymer structure or surface density, revealed qualitative differences their receptor pattern recognition (**Fig 3C**). This approach similarly highlights changes in receptor pattern recognition by different IAV strains (e.g. H1N1 vs H3N2, **Fig 5A**) or the same virus propagated in different hosts (e.g., embryonated chicken eggs or MDCK cells). Albeit, in the latter case, such differences are significantly more challenging to discern visually.

**Figure 5.**
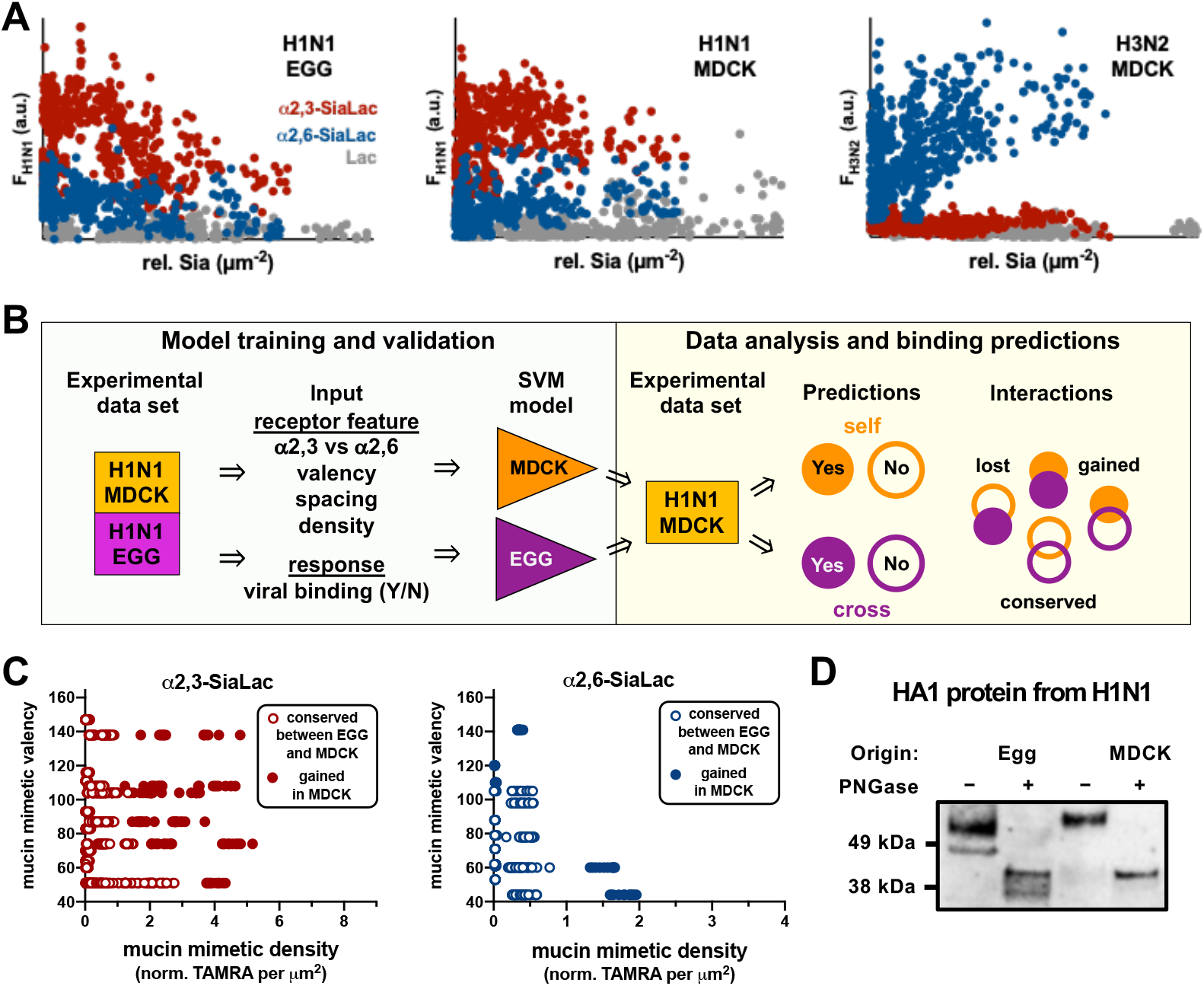
Analysis of changes in receptor pattern recognition by IAV viruses produced in avian and mammalian cells. A) Scatterplots of binding responses to changing glycan receptor density in mucin mimetic arrays for H1N1 EGG, H1N1 MDCK and H3N2 MDCK viruses. B) Experimental flow for SVM analysis of receptor pattern recognition by H1N1 viruses produced in embryonated chicken eggs and MDCK cells. Viral binding responses and receptor display parameters (i.e., glycan type, valency and spacing on mucin mimetic, mucin mimetic density and concentration in printing buffer) were used to train SVM models for each individual virus. Both models were applied to the H1N1 MDCK virus binding data set to generate predictions for binding interactions that are conserved, gained and lost as the virus is transitioning from eggs to mammalian cells. C) Based on the SVM analysis, H1N1 MDCK virus showed lower sensitivity to crowding of mucin mimetics carrying **α2**,**3-SiaLac** receptors and better utilization of **α2**,**6-SiaLac** glycans in higher valency glycoconjugates. D) Western blot analysis of HA proteins from H1N1 EGG and H1N1 MDCK viruses before and after PNGase treatment to remove *N*-linked glycans.

To aid with array data analysis, we have developed a support vector machine (SVM) approach to rapidly identify differences in receptor pattern recognition for H1N1 based on the method of their propagation (**Fig 5B** and **Fig S15**). We have used SVM to establish glycan display features to which H1N1 virus binding is conserved regardless of its origin and to identify new interactions arising as the virus adapts from eggs to mammalian cell culture. First, we established a threshold for H1N1 binding based off the background signal from the control **Lac** polymers (**Fig S16**). Then, we randomly selected a portion (67 %) of the array binding data for H1N1 EGG to train an SVM using a 5-dimensional parameter space of glycan type, glycan valency, glycan spacing on polymer, glycan density on the array, and polymer printing concentration. In combination, these parameters defined additional features of our mucin mimetic receptor displays, such as glycopolymer length (via glycan valency and glycan spacing on polymer) and glycopolymer density on the array surface (via glycan valency, glycan spacing on polymer, and glycan density on array). We used the remaining portion of the binding data set (33 %) to test accuracy, recall, and precision of the model for predicting binding interactions for H1N1 EGG in the array (**Fig S17A** and **Fig S18A**). In parallel, we trained and validated a model for H1N1 MDCK. The H1N1 EGG and H1N1 MDCK models were able to accurately produce “self-predictions” of binding and non-binding events for each respective virus in the mucin mimetic array (**Fig S17B** and **Fig S18B**).

Next, the EGG model was applied to the array binding data generated with H1N1 MDCK cells. The EGG-to-MDCK “cross-prediction” results were compared to those for “self-prediction” by the H1N1 MDCK model only (**Fig. 5B**). Within statistical error (~20%, **Fig S17** and **Fig S18**), the binding events correctly predicted by both models were termed “conserved in MDCK”. Those refer to receptor presentations that are recognized by H1N1 regardless of whether it was produced in avian or mammalian cells (**Fig S19**). The interactions correctly predicted as binding by the H1N1 MDCK model but were deemed non-binding by the H1N1 EGG model; i.e., interactions that were absent in the EGG-to-MDCK “cross-prediction”, were termed “gained in MDCK”. These refer to interactions with glycan receptor patterns in the array that were absent in the H1N1 virus produced in eggs but emerged when the virus was propagated in MDCK cells. Finally, the non-binders that were correctly predicted by the H1N1 MDCK model but were predicted to be binders by the H1N1 EGG model; i.e., interactions that were absent in H1N1 MDCK but were anticipated to occur based on the EGG-to-MDCK prediction, were termed “lost in MDCK”. These refer to glycan patterns recognized by the H1N1 virus produced in eggs but lost in mammalian cell culture.

As shown in **Figure 5C**, the conserved and gained interactions predicted by the SVM algorithm occur in distinct clusters with most of the interaction gain happening at higher polymer valencies or densities. The predictions are in agreement with our agglutination inhibition data and preliminary manual array analysis (**Fig 4** and **Fig S12**) pointing to improved ability of the H1N1 MDCK virus to engage mucin mimetic displays, both as individual polymers in solution as well as in ensembles on arrays. For **α2**,**6-SiaLac** glycopolymers, sensitivity of the virus to receptor crowding continued to persist in H1N1 MDCK, as the majority of binding gains resulted either from higher valency polymers grafted at low densities or from surface crowding of low valency mimetics (**Fig 5C**). Likewise, the SVM analysis identified stronger interactions of H1N1 MDCK with high densities of **α2**,**3-SiaLac** receptors in the mucin-mimetic array (**Fig 5C**). The predictions did not identify loss of any interactions present in the H1N1 EGG upon transition into the mammalian culture system (**Appendices 2** and **3**).

We note that an analysis of conservation, gain or loss of binding is made possible by application of the 5-dimensional SVM, wherein the algorithm learns the “rules-of-binding” from a set of interaction patterns (defined by glycan valency, density, polymer spacing and printing concentration data), which is then compared to another dataset to bring out the similarities and differences in the interactions. Arriving at these conclusions would be impossible using only chemical intuition or simple visualization of data because the interaction patterns are not apparent when the data is plotted with equal weights on all five dimensions. Using between 200 to 2000 iterations, the SVM finds linear combinations of the parameters in this 5-dimensional space, reweighting the original dimensions, until most separated classes are created, and their underlying data features are segregated, to distinguish between binding and non-binding events. It is unlikely that such combinations can be found manually, and traditional schemes, such as K-means or K-medoid, can only cluster the data and not classify them based on a target characteristic, such as binding avidity, making SVM a natural choice to create an “intelligent” space that correlates binding rules to linear combinations of glycan valency and spacing on the mimetics and their density on the array surface.

The improved range of binding interactions of H1N1 MDCK with α2,6-linked sialic acids is expected based on the higher expression of these glycans in the mammalian cells, even in absence of *Muc* gene expression in MDCK cells, the observed changes in receptor pattern recognition for both glycans are not stemming from altered affinity of its HA toward the monovalent glycans in solution (**Fig 4B**). The HA protein carries several glycosylation sites near the sialic acid binding region and the addition of glycans can influence binding^13^ and may give rise to the observed changes in receptor pattern recognition.

We performed PAGE analysis of the viral proteins before and after PNGase treatment to remove *N*-linked glycans. The glycosylated form of HA1 fragment from the egg-grown H1N1 virus had lower molecular weight than that of the MDCK virus **(Fig 5D** and **Fig S20**). This difference in gel mobility was eliminated by PNGase treatment, which catalyzes the removal of *N*-linked glycans. This points to a more extensive glycosylation of the MDCK cell-derived virus HA proteins and is consistent with previously observed differences in the glycosylation, but not in the primary amino acid sequence, of HAs from isolated duck H1N1 viruses propagated in MDCK cells and egg cultures.^43^ Whether the relationship between the changes in HA glycosylation and the altered receptor patter recognition of the virus observed in the current array study is causative or correlative is yet to be determined. Nonetheless, it presents one possible explanation for the altered binding behavior of the viruses. Such differences in glycosylation, may reflect the influence of species, cell type or combination of both on receptor pattern recognition by viruses and contribute to their emerging ability to cross between species.

## COLNCLUSIONS

The glycocalyx exists as a complex assortment of membrane-tethered and secreted glycoconjugates that serve different roles in multicellular identity, function and pathogen invasion. In this work, we aimed to model the interactions of IAVs with the mucin glycoprotein components of the mucosal barrier by generating mucin-mimetic glycopolymers with tunable sizes and compositions displaying α2,3- and α2,6-linked sialyllactose glycans as prototypes for the avian and human receptors for IAVs. RBC hemagglutination inhibition assays with soluble forms of the probes revealed an enhancement in selectivity of H1N1 PR8 viruses produced in embryonated chicken eggs from ~ 4-fold preference for binding of **α2**,**3-SiaLac** in its soluble monomeric form to more than 60-fold for the polyvalent receptor displays. This differential arose from selectively disfavored binding of the virus to the **α2**,**6-SiaLac** glycans in the mucin-mimetics rather than increased avidity toward the **α2**,**3-SiaLac** polymers. Systematic evaluation of H1N1 EGG capture on arrays of immobilized mucin mimetics enabled by an SVM learning algorithm showed that the virus bound surface displays of both receptor types but was attenuated at high polymer densities. The receptor pattern recognition changed when the virus was propagated in MDCK cells toward improved utilization of **α2**,**6-SiaLac** mucin mimetics in both soluble and immobilized forms, and a lower sensitivity to surface crowding of the **α2**,**3-SiaLac** glycopolymers.

Our findings are consistent with the proposed protective functions of soluble mucins at the airway epithelium which present primarily α2,3-linked receptors and prevent infection through viral capture and clearance. Newly, our observations that the presentation of α2,6-linked sialoglycans on linear polyvalent scaffolds and the arrangement of such conjugates in increasingly dense surface ensembles disfavors the binding of H1N1 produced in avian cells also provides support for the role of membrane-associated mucins in limiting viral adhesion and infection at the epithelial cell surface. The improved binding of H1N1 viruses produced in mammalian cells to the mucin-mimetic displays of α2,6-linked sialoglycans was not accompanied by changes in the affinity or selectivity of their HA proteins toward the individual glycan receptors. While the basis for the differences in receptor pattern recognition needs to be further investigated, our studies show that the mucin probes and their arrays may serve as useful tools to investigate viral interactions at the mucosal barrier and the evolution of their glycan receptor-binding phenotype.

## MATERIALS AND METHODS

A detailed list of chemical and biological reagents including their sources and catalog numbers can be found as Table S1 in the Supporting Information.

### Glycopolymer synthesis and characterization

Fluorescently labeled glycopolymers end-functionalized with an azide (GP_A_) used in this study and their polymeric precursors (**P**) were prepared using the RAFT polymerization method according to previously published procedures^44^ and are summarized in Figure 2 and described in detail in SI.

### Cell culture

*Madin-Darby Canine Kidney (MDCK) cells:* MDCK cells were cultured in Dulbeco’s modified Eagle’s medium supplemented with 10% FBS, 100 U/mL penicillin, and 100 U/ml streptomycin.

### Viral culture

Influenza virus strain A/PR/8/34 (H1N1, ATCC VR-1469) was purchased from ATCC and propagated in MDCK cells that were transferred to DMEM medium supplemented with 0.2% BSA fraction V, 25mM HEPES buffer, 2 µg/ml TPCK-trypsin, and 1% penicillin/streptomycin (“DMEM-TPCK” media).

The same strain was used for viral production in embryonated chicken eggs. Briefly, fertilized chicken eggs were obtained and stored at 37 °C. When the embryos were 10 days old (assessed by candeling), virus was injected through small holes made in the shell over the air sac. The holes were covered with parafilm and the eggs were placed back at 37 °C. After two days, the eggs were chilled to prepare for harvest. The eggshells were cut open above the airsac and the allantoic fluid was carefully collected into centrifuge tubes without rupturing the yolk. The tubes were centrifuged to pellet any debris and the supernatant containing virus was aliquoted into 1 mL cryovials and stored at −80 °C.

### Viral titers

Turkey red blood cells were purchased from Lampire and a 1% solution was used to determine viral titers via the hemagglutination test. MDCK cells were used to determine the 50% tissue culture infective dose (TCID50) using the Spearman-Karber method.

### Hemagglutination Inhibition

Glycopolymer solutions in PBS (25 µL, ranging from 20 µM to 20 nM by 2-fold dilutions) were added to a 96 well plate. The last well in each row was used as a PBS control and did not contain glycopolymer or virus. An equal volume (25 µL) of H1N1 diluted to HAU=4 (turkey RBCs) was added to each well and allowed to incubate at room temperature. After a ½ hr incubation, 50 µL of 1% turkey RBCs in PBS were added to all of the wells. The K_i_ value was read after a ½ hr as the lowest concentration of polymer to inhibit hemagglutination.

### Array construction

Microarrays were printed on cyclooctyne-coated glass slides as previously described^44^ using a sciFLEXARRAYER S3 printer (Scienion) following passivation with a 1% BSA/0.1% Tween-20 solution in PBS for 1 hour. Polymer solutions were diluted in printing buffer (0.005% Tween-20 in PBS) to concentrations of 1, 5 and 10 µM polymer and printed in replicates of six at a humidity of 70%. Following an overnight reaction at 4°C, excess polymer was removed by vigorous washing in 0.1% Triton X/PBS solution. The slides were then imaged on an Axon GenePix4000B scanner (Molecular Devices) at the highest PMT possible without saturation of pixels.

### Array binding assays

Prior to array binding assays, subarrays were blocked with 3% BSA solution in PBS for 1 hr. *For lectin binding*, subarrays were washed three times with lectin binding buffer (0.005% Tween-20 in PBS with 0.1 mM CaCl_2_, MnCl_2_, and MgCl_2_). Dylight labeled SNA and WGA were diluted in the lectin binding buffer and incubated on the array for 1 hour in the dark. After washing with binding buffer, 0.1% Tween-20 solution in PBS, and rinsing with MilliQ water, the slides were imaged at the highest PMT possible without producing saturated pixels. *For H1N1 binding*, subarrays were washed with 1% BSA/PBST three times following passivation. H1N1 was diluted in 1% BSA/PBST and incubated on the array for 1 hr. The slide was washed two times with 1% BSA/PBST, and then fixed for 20 min with 4 % PFA in PBS. To visualize H1N1, binding a 1:500 dilution of anti-HA in 1% BSA/PBST was incubated on the array for 1 hour, followed by an hour incubation in the dark of a 1:500 dilution of anti-rabbit-AF647 antibody. The subarrays were washed two more times with 1% BSA/PBST, two times with the 0.1% Tween-20 solution in PBS, rinsed with MilliQ and imaged at the highest PMT possible without producing saturated pixels.

### Machine learning workflow

Each experiment containing triplicates of fluorescence measurement of 6 samples (18 measurements per experiments) were treated independently. The workflow is demonstrated in Figure S15. The fluorescence intensities were normalized over the entire data set in the range of [0,1]. This continuous data was converted to categorical (“bound”/”not bound”) based on a cut-off fluorescence that was determined from the distribution of fluorescence intensities of lactose samples which served as negative control. Next, the features (Glycan type, Valency, and polymer density) were scaled to a range of [0,1] in order to avoid bias from higher values. The only categorical feature (glycan type) was transformed into numeric by mapping to a two-dimensional space were 2,3-SiaLac is represented by (1,0) and 2,6-SiaLac by (0,1). This data set was then split into a training and test set, where training set contained 66% of the data. Each experiment of 18 measurements was individually split between training and test. Support Vector Machine (SVM) algorithm was used for learning from the training data, and predictions were validated against the test data. Training and testing were performed separately for egg-virus data and MDCK-virus data. Convergence was reached within 2000 iterations for egg virus data and 200 iterations for MDCK data (Fig S17). Next, the algorithm trained on egg-virus data was used to predict results for MDCK-virus data, and the prediction results from this so-called “cross-model” was compared with the results obtained from the model trained on MDCK-virus (the “self-model”). Data was prepared in python using pandas, numpy, and scipy packages. SVM was performed using python’s scikit-learn package.^45^

## SUPPORTING INFORMATION

General procedures for glycopolymer synthesis, HI assays, microarray construction, lectin and H1N1 binding to arrays, and machine learning algorithm design details can be found in the Supporting Information. Additional controls for HI assays and fluorescent scans of the glycan microarrays with lectins (SNA and WGA) and virus bound can also be found in the Supporting Information. The NMRs used to characterize the glycopolymers can be found in Appendix 1.

## FUNDING SOURCES

This work was supported in part by the NIH Director’s New Innovator Award (NICHD: 1DP2HD087954-01), the NIH Director’s Glycoscience Common Fund (1R21AI129894-01) and the Gordon and Betty Moore Foundation via a Scialog grant (#9162.07). KG was supported by the Alfred P. Sloan Foundation (FG-2017-9094) and the Research Corporation for Science Advancement via the Cottrell Scholar Award (grant #24119). TML was supported by the Chemistry-Biology Interface training program (NIGMS: 5T32GM112584-03) and CJF was supported by the Molecular Biophysics Training Program (T32 GM08326). PG and MA are supported by the G. Harold and Leyla Y. Mathers Charitable Foundation.

## ACKNOWLEDGMENTS

We thank the UCSD Glycobiology Research and Training Center for access to tissue culture facilities and analytical instrumentation, Dr. Christopher Fisher for his help with virus culture and characterization methods, and the ASU Research Computing and their Agave computing cluster for the computational resources. The authors also wish to thank Prof. Mia Huang (TSRI) for her technical advice and valuable insights over the course of this research.

## Notes

### Competing Interest Statement

The authors have declared no competing interest.

