## Supporting information for "Mucin-mimetic glycan arrays integrating machine learning for analyzing receptor pattern recognition by influenza A viruses"

**Abbreviations**

IAV = influenza A virus
HA = hemagglutinin
NA = neuraminidase
GPB = glycan binding protein
RBC = red blood cell
MDCK = Madin-Darby canine kidney
BSA = bovine serum albumin
PBS = phosphate buffered saline
HI = hemagglutination inhibition
HAU = hemagglutination units
$TCID_{50}$ = tissue culture infective dose (50%)
MOI = multiplicity of infection
SA = sialic acid
HPLC = high performance liquid chromatography
RAFT = reversible addition-fragmentation chain transfer
PBST = PBS with 0.05% Tween-20
DBCO = dibenzocyclooctyne
DP = degree of polymerization
MW = molecular weight
Đ = dispersity
CTA = chain transfer agent
GPC = gel permeation chromatography
SEC = size exclusion chromatography
TAMRA = tetramethylrhodamine
AF647 = AlexaFluor 647
Boc = ditertbutylcarbonate
SPAAC = strain promoted azide-alkyne cycloaddition
$K_i$ = inhibitory constant
SNA = *sambucus nigra* agglutinin
WGA = wheat germ agglutinin
LBB = lectin binding buffer

**Table S1. Chemical and biological reagents.**

| Reagent | Source | Cat. No. |
|---|---|---|
| 3'-Sialyllactose sodium salt | Carbosynth | OS04397 |
| 6'-Sialyllactose sodium salt | Carbosynth | OS04398 |
| Amicon Ultra-0.5 mL 10K Spin filters | Millipore | UFC5010 |
| Dibenzocyclooctyne (DBCO) amine | BroadPharm | BP-22066 |
| SuperChip Microarray Slides Epoxysilane | Thermo Fisher | EP10143261 |
| TAMRA Maleimide | Thermo Fisher | T6027 |
| BSA Fraction V | Spectrum | A3611 |
| Dylight649 conjugated *Sambucus nigra* lectin | Ey Labs | DY649-6802-1 |
| Dylight649 conjugated *Wheat germ agglutinin* lectin | Ey Labs | DY649-2101-1 |
| Influenza virus A/PR/8/34 (H1N1) | ATCC | VR-1469 |
| PBS (with Calcium and Magnesium) | Gibco | 14040141 |
| PCR tube | Thermo Fisher | AB0337 |
| PD-10 Desalting Column | GE Life Sciences | 17085101 |
| Turkey Red Blood Cells | Lampire | 7249408 |
| Phosphate buffered saline | Gibco | 10010023 |
| A/California/06/09 H1N1 (polyclonal rabbit IgG α-HA antibody) | eEnzyme | IA-01SW-0100 |
| goat α-rabbit-AF647 | Thermo Fisher | A21244 |
| Bolt 4-12% Bis-Tris Plus gel | Invitrogen | NW04120BOX |

| Reagent | Source | Cat. No. |
|---|---|---|
| PNGase F | New England Biolabs | P0704L |
| Anti-rabbit IgG, HRP-linked Antibody | Cell Signaling Technologies | 7074s |
| SeeBlue Plus2 Prestained standard ladder | Invitrogen | LC5925 |
| BOLT LDS sample Buffer (4x stock) | Invitrogen | B0007 |
| NuPAGE™ MES SDS Running Buffer (20X) | Invitrogen | NP0002 |
| iBlot2 PVDF Mini Stacks | Invitrogen | IB24002 |
| Luminata Classico HRP substrate | Millipore | WBLUC0500 |

*All other chemicals, if not otherwise indicated, were purchased from Sigma-Aldrich.

### Instrumentation

Proton nuclear magnetic resonance ($^1$H NMR) spectrum of polymer backbones and glycopolymers were obtained on either a 300 MHz (Bruker) or 500 MHz (Joel) NMR spectrometer, using deuterated solvents ($CDCl_3$ for the polyacrylaminde precursors, **P**, and sodium phosphate buffered saline (150 mM) in $D_2O$ adjusted to pH 7.4 with DCl for the glycopolymers). The spectra were analyzed using MestReNova software and are reported in parts per million (ppm) on the δ scale relative to the residual solvent as an internal standard (for $^1$H NMR: $CDCl_3$ = 7.26 ppm, $D_2O$ = 4.79 ppm). Data are reported as follows: chemical shift, multiplicity (s = singlet, d = doublet, dd = doublet of doublets, t = triplet, q = quartet, br = broad, m = multiplet), and integration.

Size exclusion chromatography was performed on a Hitachi Chromaster system equipped with an RI detector and a 5 µm, mixed bed, 7.8 mm I.D. x 30 cm TSKgel column (Tosoh Bioscience). Polymer backbones **S/M/L-P** were analyzed in DMF (0.2% w/v LiBr, 70°C) using an isocratic method with a flow rate of 0.7 mL/min. Glycan conjugation reactions were performed in a Biorad MyCycler thermocycler (Hercules, CA).

Glycan microarrays were printed on cyclooctyne-coated glass slides as previously described[1] using a sciFLEXARRAYER S3 printer (Scienion) and imaged on an Axon

GenePix4000B scanner (Molecular Devices) at the highest PMT possible without saturation of pixels.

Protein gels were run using the Invitrogen Mini Gel Tank (Cat no. A25977) set at constant 200 V for 22 minutes and transferred to PVDF membranes using the Invitrogen iBlot2 transfer system (Cat. no. IB21001) with P3 settings (20 V for 7 min). Following probing with antibodies, the blots were imaged using a BioRad GelDoc XRS+ imaging system

**Preparation and characterization of mucin mimetic glycopolymers GP.**

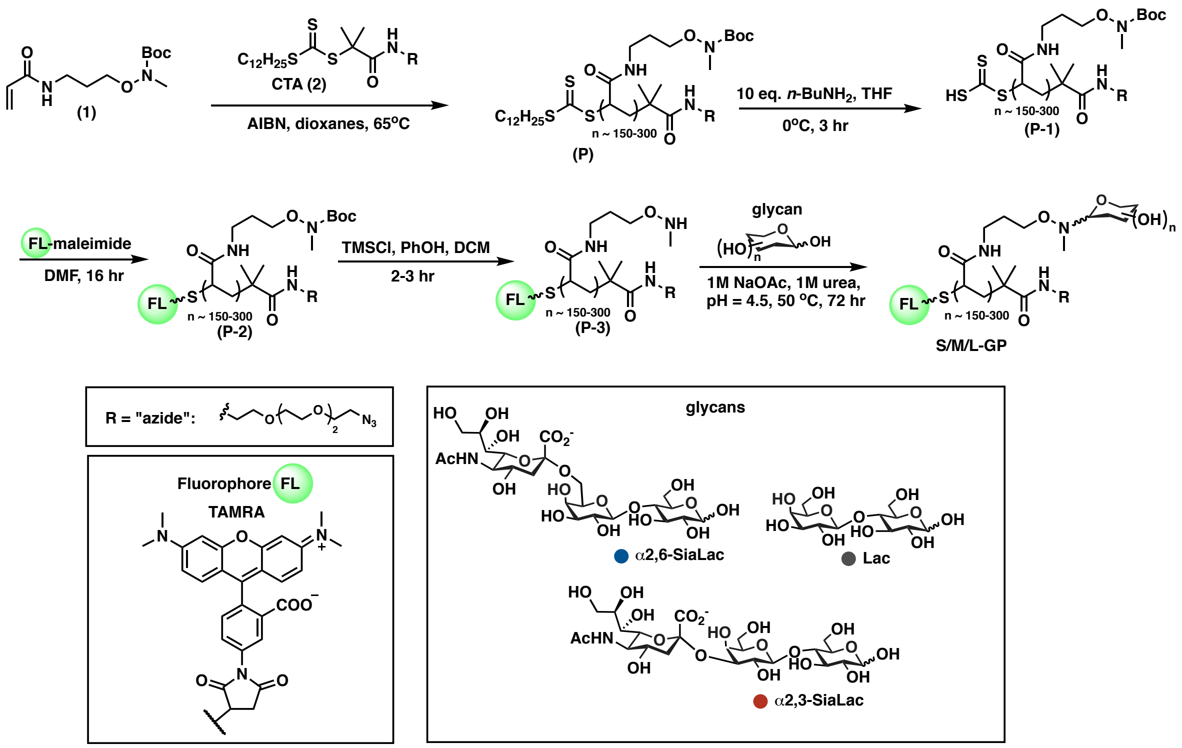

**Scheme S1. Preparation of mucin mimetic glycopolymers GP**

**Preparation of polyacrylamide precursors (P).** A 10 mL flame-dried Schlenk flask equipped with a magnetic stir bar was charged with monomer **1** and CTA **2** according to ratios listed in Table S2. Dry dioxane was added, followed by an aliquot of AIBN stock solution (2 mg/mL, Table S2).

The reaction volume was adjusted with dry dioxane to a final monomer concentration of 1.5 M. The yellow solution was thoroughly degassed through at least three freeze-pump-thaw cycles and backfilled with $N_2$. The flask was then heated to 65˚C and allowed to react for 8-12 hrs as listed in Table S2. The reaction was stopped at ~ 60-70% conversion by precipitation into hexanes. The resulting yellow solid was dissolved in DCM and precipitated again into hexanes. This purification step was repeated twice. The final polymer was concentrated from $CHCl_3$ to remove residual hexanes and dried under vacuum to give polymers **P** as pale-yellow solids. To achieve different polymer lengths, the monomer to CTA ratio was adjusted based on the theoretical degree of polymerization (DP) and assuming 70% consumption of monomer (see Table S2). All **P** polymers were analyzed by SEC in DMF.

**Table S2. Polymer backbone characteristics**

| Polymer ID | Mole ratio M(1)/CTA(2) | 1 Monomer (mmol) | 2 CTA (µmol) | AIBN (mmol) | Rxn time (hr) | % yield | Mw (kDa) | DP via SEC | *Đ* |
|---|---|---|---|---|---|---|---|---|---|
| **S-P** | 200 | 2.57 | 12.9 | 2.56 | 8 | 90 | 39 | 150 | 1.28 |
| **M-P** | 300 | 1.94 | 6.47 | 1.28 | 10 | 87 | 54 | 200 | 1.17 |
| **L-P** | 600 | 3.59 | 6.03 | 2.44 | 12 | 91 | 98 | 300 | 1.36 |

**Preparation of chain end-deprotected polymer precursors (P-1).** The polymer backbones **P** were dissolved in anhydrous THF to give a 2 mM solution. Next, a solution of n-butylamine in THF (20 mM, 10 equiv. per trithiocarbonate end group) was added and stirred on ice for 2-3 hr. The product was precipitated in excess hexanes three times before concentrating in $CHCl_3$ and drying to give polymers as a white solid (yield: 85-95% for **P-1**).

**Preparation of TAMRA labeled polymer precursors (P-2).** This step was carried out immediately after end deprotection. A 2 mM solution of fluorophore-maleimide (1.5 eq., TAMRA per polymer end group) in dry DMF was used to dissolve end deprotected polymers (**P-1)** in a 4 mL vial equipped with a stir bar. The reactions were stirred in the dark overnight. The reaction mixtures were then diluted in ether before being precipitated in excess hexanes to remove unreacted fluorophore. The mixtures were centrifuged (1000 xg, 3 min) and the hexanes was decanted and replaced with fresh hexanes so the precipitation could be repeated. Finally, the products were concentrated using $CHCl_3$, and dried under vacuum to give a dark pink solid (**P-2**). (Yields: 94-96% for **P-2**)

**Preparation of side chain deprotected polymer precursors (P-3).** The TAMRA labeled polymers were charged into 4 mL vials and dissolved in 500 μL of 1 M TMS-Cl and 3 M phenol mixture freshly prepared in anhydrous DCM. The solutions were stirred for 2 hr in the dark. The product was precipitated into ether three times from DCM, redissolved in MilliQ water and purified on a PD10 column. The product was the lyophilized overnight to give polymers (**P-3**) as a fluffy pink solid (**P-3**). (Yields: 70-75% for **P-3**)

**Preparation of mucin mimetic glycopolymers GP.** The lyophilized deprotected polymers were dissolved in sugar ligation buffer (1M NaOAc, 1M urea, pH 4.5) to achieve a solution where the side chain concentration was kept constant at 200 mM. The solutions were then added to PCR tubes containing 0.1 to 1.1 equivalents of **α2,3-SiaLac**, **α2,6-SiaLac**, or **Lac** according to Table S3 and heated at 50˚C for 72 hr on a thermocycler. These reactions were purified using a Amicon Ultra Centrifugal Filter (pre-washed with Milli-Q water twice by spin dialysis, 10K MWCO, Millipore) and spin dialyzed (10000 xg, 15 min) five times using 500 μL of deuterated phosphate buffered saline solution (100 mM phosphate, 150 mM NaCl, pD 7.4). The glycan ligation efficiency was determined from the $^{1}$H-NMR spectra of the glycopolymers by subtracting polymer backbone protons (total of 7 protons, found between 2.5-4.5 ppm) from the total integration in the region 2.5-4.5 ppm and dividing the difference by the number of glycan protons which are also found between 2.5-4.5 ppm. Glycan valency was then calculated as the product of the degree of polymerization and ligation efficiency. To determine glycopolymer labeling efficiency, UV-Vis experiments were performed using a quartz cuvette (10 mm path length) in a Nanodrop2000c spectrophotometer (ThermoFisher). The TAMRA absorbance at a wavelength of 555 nm was measured for a known concentration (by weight) of glycopolymer. The fluorophore concentration of the solution was obtained using Beer's Law, and the fluorophore labeling efficiency was calculated by dividing the fluorophore concentration by the known polymer concentration. This value, which was determined for each polymer backbone, was then used to adjust the polymer concentration obtained in the UV-Vis measurements. All of the glycopolymer characteristics are displayed in Table S3.

**Table S3. Structural characteristics of glycopolymers GP.**

| Entry # | Polymer ID | DP | TAMRA labeling (%) | Glycan | Glycan equiv. added | Glycan ligation efficiency (%) | Glycan valency | Theoretical end-to-end length (nm) |
|---|---|---|---|---|---|---|---|---|
| **1** | **S-$^{3}$GP$_{50}$** | 150 | 6.3 | **α2,3 SiaLac** | 0.49 | 34 | 50 | 8 |
| **2** | **S-$^{3}$GP$_{60}$** | 150 | 27.6 | **α2,3 SiaLac** | 1.10 | 40 | 60 | 8 |
| **3** | **S-$^{3}$GP$_{85}$** | 150 | 6.3 | **α2,3 SiaLac** | 1.09 | 58 | 85 | 8 |
| **4** | **M-$^{3}$GP$_{50}$** | 200 | 40.8 | **α2,3 SiaLac** | 0.09 | 27 | 50 | 9 |
| **5** | **M-$^{3}$GP$_{65}$** | 200 | 40.8 | **α2,3 SiaLac** | 0.25 | 34 | 65 | 9 |

| | | | | | | | | |
|---|---|---|---|---|---|---|---|---|
| **6** | **M-$^{3}$GP$_{70}$** | 200 | 40.8 | **α2,3 SiaLac** | 0.35 | 37 | 70 | 9 |
| **7** | **M-$^{3}$GP$_{75}$** | 200 | 27.5 | **α2,3 SiaLac** | 0.53 | 39 | 75 | 9 |
| **8** | **M-$^{3}$GP$_{80}$** | 200 | 40.8 | **α2,3 SiaLac** | 0.49 | 43 | 80 | 9 |
| **9** | **M-$^{3}$GP$_{90}$** | 200 | 40.8 | **α2,3 SiaLac** | 0.71 | 49 | 90 | 9 |
| **10** | **M-$^{3}$GP$_{110a}$** | 200 | 40.8 | **α2,3 SiaLac** | 1.08 | 58 | 110 | 9 |
| **11** | **M-$^{3}$GP$_{110b}$** | 200 | 27.5 | **α2,3 SiaLac** | 1.18 | 54 | 110 | 9 |
| **12** | **L-$^{3}$GP$_{110}$** | 300 | 9.5 | **α2,3 SiaLac** | 0.68 | 36 | 110 | 12 |
| **13** | **L-$^{3}$GP$_{140}$** | 300 | 9.5 | **α2,3 SiaLac** | 1.14 | 46 | 140 | 12 |
| **14** | **L-$^{3}$GP$_{145}$** | 300 | 55.8 | **α2,3 SiaLac** | 1.09 | 49 | 145 | 12 |
| **15** | **S-$^{6}$GP$_{45}$** | 150 | 6.3 | **α2,6 SiaLac** | 0.47 | 29 | 45 | 8 |
| **16** | **S-$^{6}$GP$_{60}$** | 150 | 27.6 | **α2,6 SiaLac** | 1.09 | 41 | 60 | 8 |
| **17** | **S-$^{6}$GP$_{65}$** | 150 | 6.3 | **α2,6 SiaLac** | 1.10 | 43 | 65 | 8 |
| **18** | **M-$^{6}$GP$_{50}$** | 200 | 40.8 | **α2,6 SiaLac** | 0.11 | 28 | 50 | 9 |
| **19** | **M-$^{6}$GP$_{60}$** | 200 | 40.8 | **α2,6 SiaLac** | 0.26 | 32 | 60 | 9 |
| **20** | **M-$^{6}$GP$_{70}$** | 200 | 40.8 | **α2,6 SiaLac** | 0.35 | 37 | 70 | 9 |
| **21** | **M-$^{6}$GP$_{80}$** | 200 | 40.8 | **α2,6 SiaLac** | 0.51 | 41 | 80 | 9 |
| **22** | **M-$^{6}$GP$_{90}$** | 200 | 40.8 | **α2,6 SiaLac** | 0.69 | 46 | 90 | 9 |
| **23** | **M-$^{6}$GP$_{100}$** | 200 | 27.5 | **α2,6 SiaLac** | 1.13 | 51 | 100 | 9 |
| **24** | **M-$^{6}$GP$_{105}$** | 200 | 40.8 | **α2,6 SiaLac** | 1.07 | 55 | 105 | 9 |
| **25** | **L-$^{6}$GP$_{105}$** | 300 | 9.5 | **α2,6 SiaLac** | 0.68 | 35 | 105 | 12 |
| **26** | **L-$^{6}$GP$_{120}$** | 300 | 37.4 | **α2,6 SiaLac** | 1.11 | 40 | 120 | 12 |
| **27** | **L-$^{6}$GP$_{140}$** | 300 | 9.5 | **α2,6 SiaLac** | 1.08 | 47 | 140 | 12 |

| | | | | | | | | |
|---|---|---|---|---|---|---|---|---|
| **28** | **S-ØGP$_{65}$** | 150 | 27.6 | **Lac** | 0.51 | 44 | 65 | 8 |
| **29** | **S-ØGP$_{75}$** | 150 | 6.3 | **Lac** | 0.49 | 51 | 75 | 8 |
| **30** | **S-ØGP$_{110}$** | 150 | 6.3 | **Lac** | 1.08 | 74 | 110 | 8 |
| **31** | **M-ØGP$_{85}$** | 200 | 40.8 | **Lac** | 0.11 | 44 | 85 | 9 |
| **32** | **M-ØGP$_{95a}$** | 200 | 27.5 | **Lac** | 0.58 | 47 | 95 | 9 |
| **33** | **M-ØGP$_{95b}$** | 200 | 40.8 | **Lac** | 0.25 | 50 | 95 | 9 |
| **34** | **M-ØGP$_{110}$** | 200 | 27.5 | **Lac** | 0.37 | 57 | 110 | 9 |
| **35** | **M-ØGP$_{130}$** | 200 | 27.5 | **Lac** | 1.22 | 66 | 130 | 9 |
| **36** | **L-ØGP$_{85}$** | 300 | 9.5 | **Lac** | 0.22 | 28 | 85 | 12 |
| **37** | **L-ØGP$_{155}$** | 300 | 37.4 | **Lac** | 0.70 | 52 | 155 | 12 |
| **38** | **L-ØGP$_{170}$** | 300 | 9.5 | **Lac** | 0.66 | 56 | 170 | 12 |

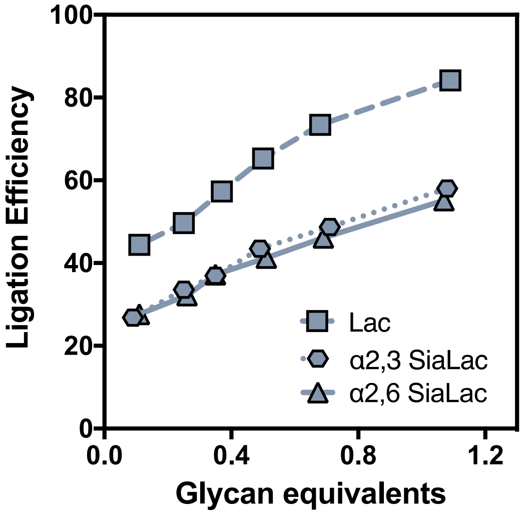

**Figure S1. Control of glycopolymer valency.** The number of glycans introduced into the polymers was controlled through stoichiometry of **SiaLac** or **Lac** with respect to the reactive side chains. The uncharged disaccharide Lac is incorporated more readily than the negatively charged **SiaLac** trisaccharides.

### Calculations for polymer end-to-end length in solution

The polymer length in solution was calculated as previously described[2] based on the theoretical distance between polymer chain termini (R), assuming a hard sphere model for atoms and imposing the excluded volume effect. The calculation is as follows:

**Equation S1.** $$R = \left(\frac{2}{3}\right)^{0.5} N^{0.6} C^{0.2} b^{0.4} \left(\frac{4\pi b^3}{3}\right)^{0.2}$$

where *N* is the number of C-C bonds between the side chain (DP x 2), *C* is the polymer characteristic ratio (8.5 for polyacrylamide),[3] and *b* is the C-C bond length (0.154 nm).

### Mucin-mimetic array preparation and analysis with SNA and WGA plant lectins

**Construction and characterization of density-variant mucin mimetic arrays.** All pre-print passivation solutions, printing buffers, and the system liquid were first filtered through 0.22 µm filters. For initial cyclooctyne functionalization, epoxysilane slides (Thermo Scientific SuperChip Microarray Slides) were incubated in a Coplin jar with rocking in a 1 mM solution of dibenzocyclooctyne (DBCO) amine (BroadPharm) in DIPEA/DMF overnight. Slides were then sonicated two times for 15 minutes in MeOH, rinsed with MilliQ, and spin-dried (550 rpm for 5 min). They were stored at 4˚C with desiccant until printing.

On the day of printing, the slides are passivated with a 1% BSA/0.1% Tween-20 solution in PBS for 1 hour with rocking, then washed with MilliQ three times for 15 min and spin-dried. A Scienion sciFLEXARRAYER S3 printer was used to print 6 replicates of polymers diluted in printing buffer (0.005% Tween-20 in PBS) at a humidity of 70%. After printing, the slides were re-humidified over an 80˚C water bath, snap dried on a glass plate set at 80˚C, and allowed to react at 4˚C overnight.

After outlining the subarrays with a glass cutter and snap drying again at 80˚C, the slides were immediately washed vigorously in a 0.1% TritonX/PBS solution for 2 minutes and rocked for another 15 minutes. The slides were next washed in PBS two times for 10 minutes, rinsed with MilliQ, and spin-dried. The slides can be stored at 4˚C with desiccant until further use.

**Lectin binding in density variant mucin-mimetic arrays.** For lectin binding, the slides were washed for 15 minutes in a 0.1% Tween-20/PBS solution on a shaker and then spin-dried. They were then imaged at the highest PMT possible without saturation using an Axon GenePix 4000B microarray scanner (Molecular Devices). A gasket was added, and the subarrays were blocked with 3% BSA solution in PBS for 1 hour at RT, then washed three times with lectin binding buffer (LBB: 0.005% Tween-20 in PBS with 0.1 mM $CaCl_2$, $MnCl_2$, and $MgCl_2$). Dylight labeled SNA and WGA was diluted in the LBB and incubated on the array for 1 hour at room temperature with rocking and in the dark. The labeled lectin is removed, and the subarrays are washed two times with LBB, two times with the 0.1% Tween-20 solution, rinsed with MilliQ and spin-dried. The slides were imaged at the highest PMT possible without producing saturated pixels.

**Array image collection and processing.** GenePix Pro v7 software was used to image and analyze the microarrays. A block was constructed with the same dimensions as the printed array, aligned over the spots in the image, and analyzed. To determine the amount of polymer immobilized on the slide, the mean background subtracted 532 nm signal was used. Dividing this signal by the labeling efficiency and multiplying by the polymer valency provided the amount of relative glycan per spot (Fig S2). The spot area was calculated by using the spot diameter from the generated results page. Dividing the relative glycan amount by the spot area provided the relative glycan density per μm$^2$. The signal at 635 nm after background subtraction was used to analyze binding of Dylight649-SNA and WGA or H1N1 PR8 probed with AF647-anti-HA Ab. For viral binding experiments, an AF647 anti-HA antibody only staining was used to determine background.

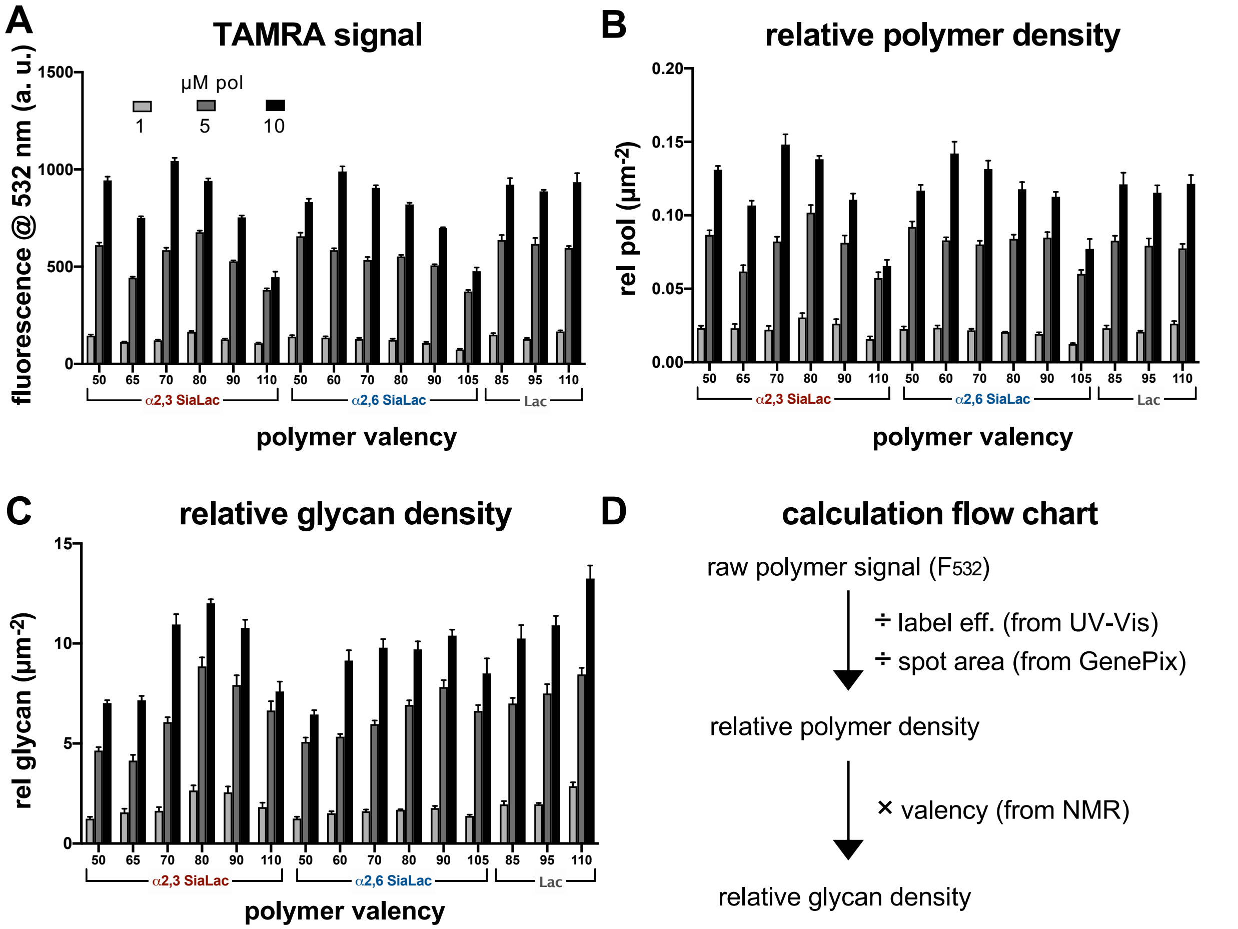

**Figure S2. Conversion of TAMRA signal to relative polymer density and relative glycan density for the M-GP array.** (A) The TAMRA signal of the labeled polymers is obtained from analysis using GenePix software. (B) The TAMRA signal can be converted to relative polymer density by dividing by the polymer labeling efficiency (41%) acquired through UV-Vis spectrometry measurements and dividing by the spot area which is calculated using the diameter from GenePix analysis. (C) The relative glycan density can be determined by multiplying the relative polymer density by the valency of each polymer which is obtained through NMR integration. (D) A flowchart showing the calculations.

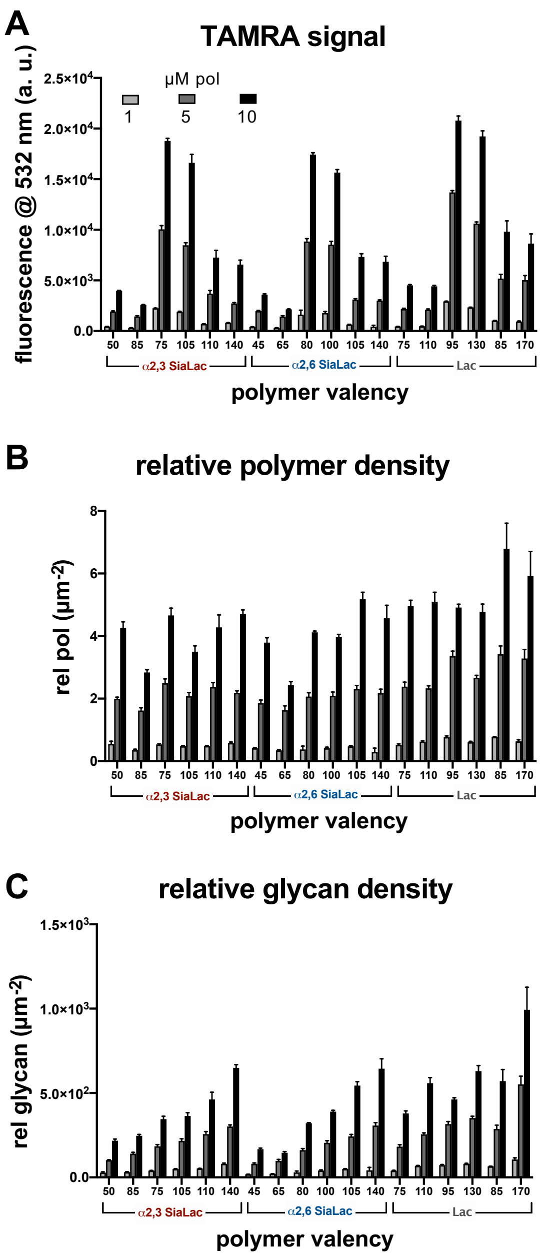

**Figure S3. Polymer grafting and glycan density on arrays containing all polymer lengths.** (A) The TAMRA signal of the labeled polymers is obtained from analysis using GenePix software. (B) The TAMRA signal can be converted to relative polymer density by dividing by the polymer labeling efficiency (S: 6%, M: 28%, L: 9%) acquired through UV-Vis spectrometry measurements and dividing by the spot area which is calculated using the diameter from GenePix analysis. (C) The relative glycan density can be determined by multiplying the relative polymer density by the valency of each polymer which is obtained through NMR integration. (D) A flowchart showing the calculations.

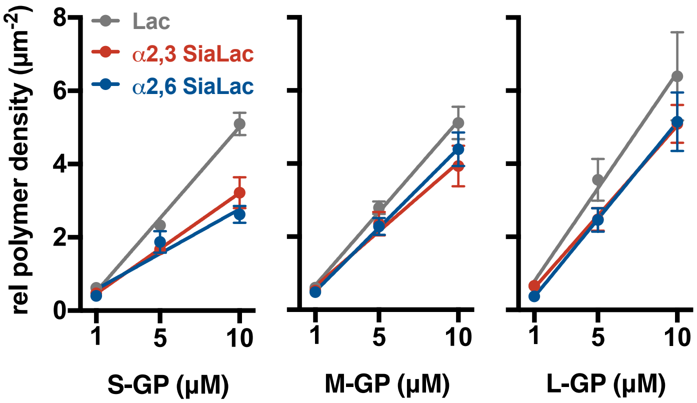

**Figure S4. Relative array surface density for polymers S-, M-, and L- GPs with maximal glycosylation.** Data points represent means ± standard deviation of 12 replicate spots and is fit using linear regression analysis (PRISM). All GPs remain in the linear range of grafting to the array surface. Lac polymers achieve the greatest density for all polymer sizes while the SiaLac polymers graft similarly under all conditions.

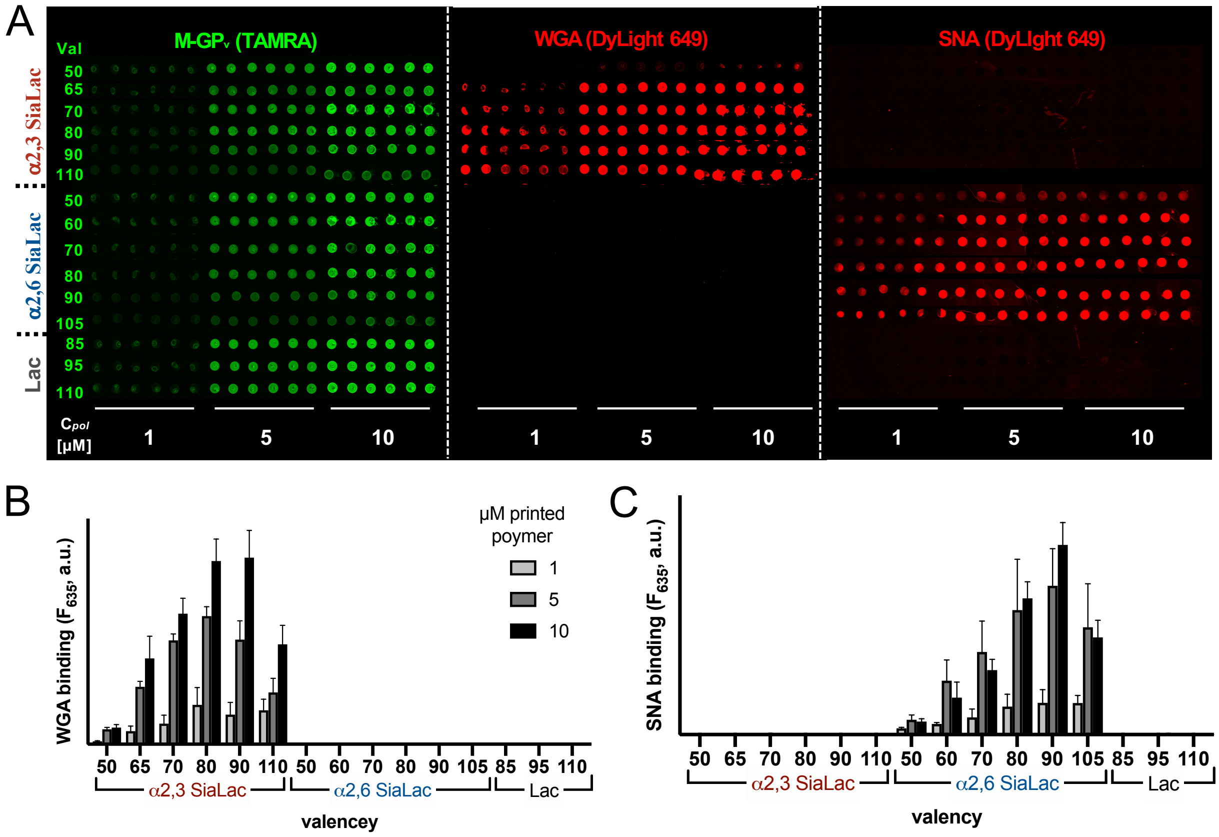

**Figure S5. SNA and WGA binding in mucin-mimetic arrays (extended data for Figure 3).** (A) Micrographs of the printed glycopolymers (TAMRA) and the bound lectins (DyLight649). WGA binds only to M-$^{3}GP_{50\text{-}110}$ (middle panel), and AF647 labeled SNA binds only to $^{6}GP_{50\text{-}105}$. Bar graph representations for (B) WGA binding at 500 nM and (C) SNA binding at 500 nM. Error bars represent standard deviation from 6 replicate spots.

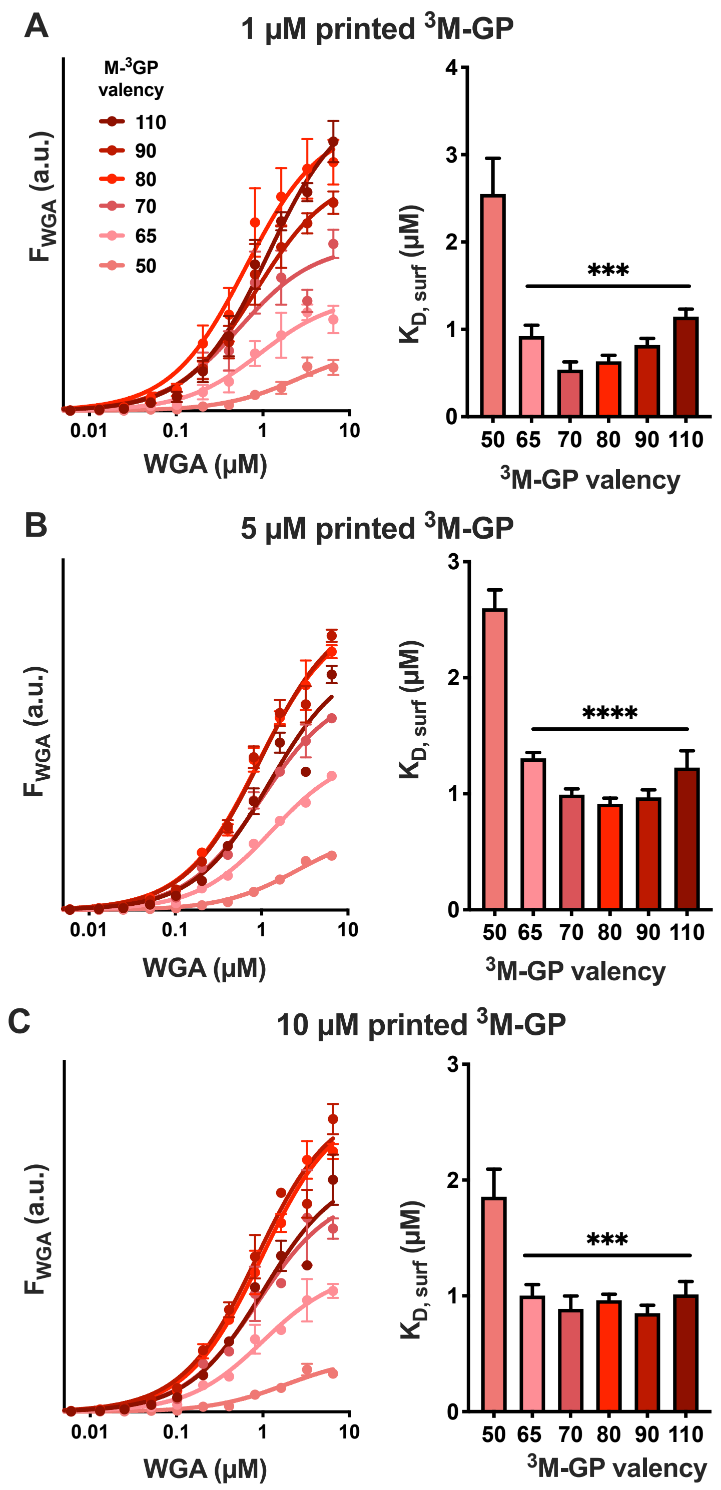

**Figure S6. Determination of KD, surf for WGA binding to M-3GP$_{50\text{-}110}$ in arrays.** Binding of WGA to the glycopolymers follows the same trends at all printing concentrations, however the valency dependence on $K_{D,\,surf}$ is most prominent at 1 µM printed polymer (A), decreases at 5 µM (B), and is essentially nonexistent for $^{3}$**M-GP$_{65\text{-}100}$** at the 10 µM printing concentration.

**Table S4. $K_{D, surf}$ values for the binding of WGA to M-$^3$GP$_{50-110}$**

| | µM printed polymer | | |
|---|---|---|---|
| **val** | **1** | **5** | **10** |
| **50** | 2.5 ± 0.4 | 2.6 ± 0.2 | 1.8 ± 0.3 |
| **65** | 0.9 ± 0.1 | 1.30 ± 0.06 | 0.98 ± 0.09 |
| **70** | 0.54 ± 0.08 | 1.01 ± 0.05 | 0.9 ± 0.1 |
| **80** | 0.63 ± 0.06 | 0.91 ± 0.05 | 0.95 ± 0.06 |
| **90** | 0.82 ± 0.07 | 0.95 ± 0.07 | 0.88 ± 0.08 |
| **110** | 1.15 ± 0.08 | 1.2 ± 0.2 | 1.1 ± 0.1 |

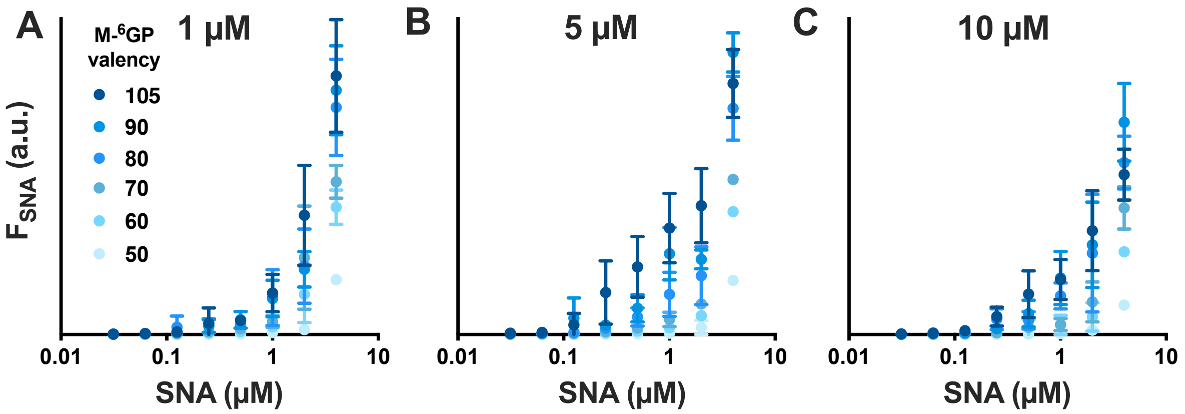

**Figure S7. Determination of KD, surf for SNA binding to M-6GP50-110 in arrays.** A-C) Fluorescence intensity of Dyight649-SNA bound to arrays printed at 1, 5 and 10 µM concentrations. Saturation binding could not be achieved due to SNA aggregation.

**Crystal structures of WGA and SNA lectins.**

The lectin cartoons generated for Fig 3C were adapted from crystal structures of WGA complexed with a bivalent sialoglycan peptide[4] (PDB ID: 2CWG) and SNA-II complexed with lactose[5] (PDB ID: 3CA4). In the cartoons in Fig 3, ovals were used to approximate the backbone of the protein from the crystal structures and the binding sites were designated as gray diamonds.

WGA exists as a dimer, and while it is most commonly known for binging GlcNAc, it can also bind sialic acid, as shown in the crystal structure. SNA-I has not been crystalized, but it is a related lectin to SNA-II that exists can exist in multiple oligomeric states (monomer, dimer, or tetramer) and is specific for α2,6-linked sialic acid.[6]

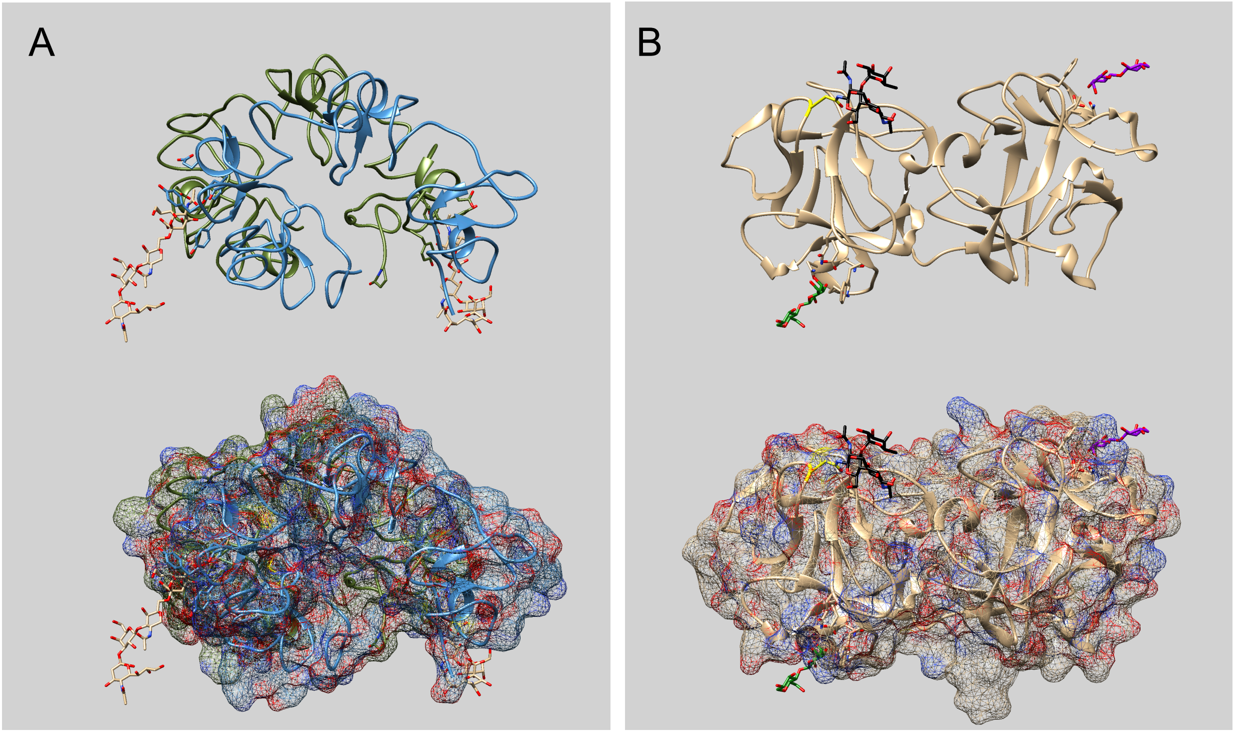

**Figure S8. Crystal structures of dimeric WGA and monomeric SNA in ribbon and mesh representations.** A) The WGA dimer is shown in complex with a sialyllated glycopeptide. This lectin contains two sialic acid binding sites on the same side of protein separated by 3.9 nm. B) Monomeric SNA contains two binding sites per subunit that are found on opposite ends of the protein. The glycan in the Iα binding site is colored in green and the one in the IIγ binding site is colored in purple. The black *N*-linked glycan included in the structure is important for the oligomerization state of the lectin.

We also performed a BLASTp analysis of the SNA-I binding site with that of SNA-II to determine the homology between the receptor binding domains.

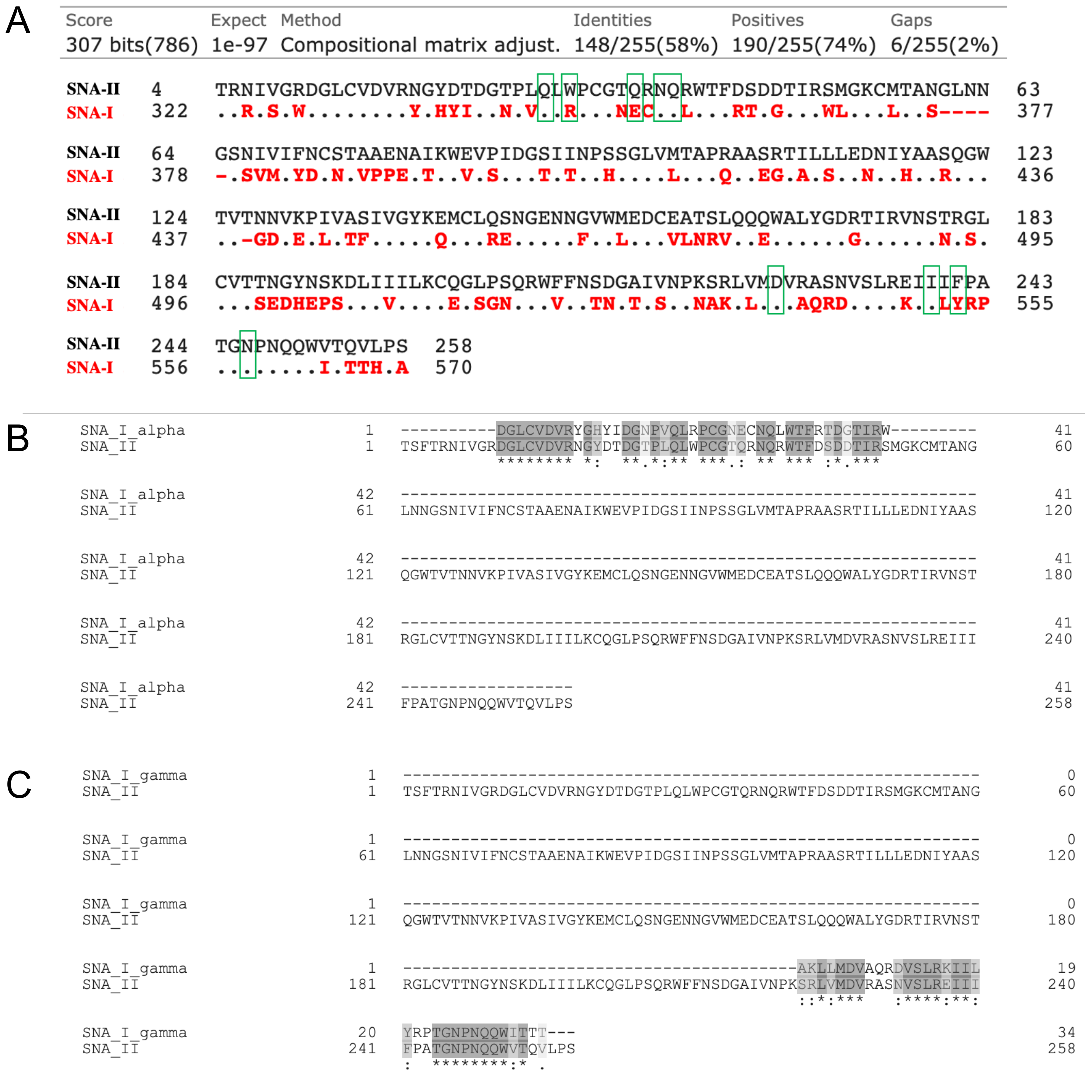

**Figure S9. BLASTp sequence alignment of SNA-I and SNA-II.** A) BLASTp alignment of the whole proteins for SNA-I and SNA-II. Amino acids that bind to the glycan receptors are outlined in green. Conserved residues are shown as dots in the SNA-I sequence. B) Local alignment of the Iα tandem repeat domain that contains the carbohydrate binding site to SNA-II. Alignment shows strong similarity to the reported Iα carbohydrate binding site of SNA-II. Dark gray boxes indicate conserved amino acids, and the light gray boxes denote amino acids of similar structure. C) Local alignment of the IIγ tandem repeat domain carbohydrate binding site to SNA-II. Alignment shows strong homology to the reported IIγ carbohydrate binding site of SNA-II.

### Inhibition of H1N1 agglutination of RBCs by free glycans and soluble glycopolymers (GPs)

**Hemagglutination inhibition (HI) assay procedure.** The viral stock solution was diluted to an HAU = 4 for HI experiments. This HAU was tested to ensure that it consistently hemagglutinated RBCs in the absence of soluble inhibitors. Glycopolymer solutions were diluted to the same starting concentration of 20 μM polymer (200 μM for the monovalent glycans) in the first lane of a 96 well plate, and a 2-fold dilution to a total volume of 25 μL was performed down the plate. The last well was used as a PBS only control. Then 25 μL of the viral dilution (HAU = 4) was added and incubated for a ½ hour at room temperature. After which time, 50 μL of a 1% turkey RBC solution was added to all wells. The assay was carried out in duplicate and read after ½ hour.

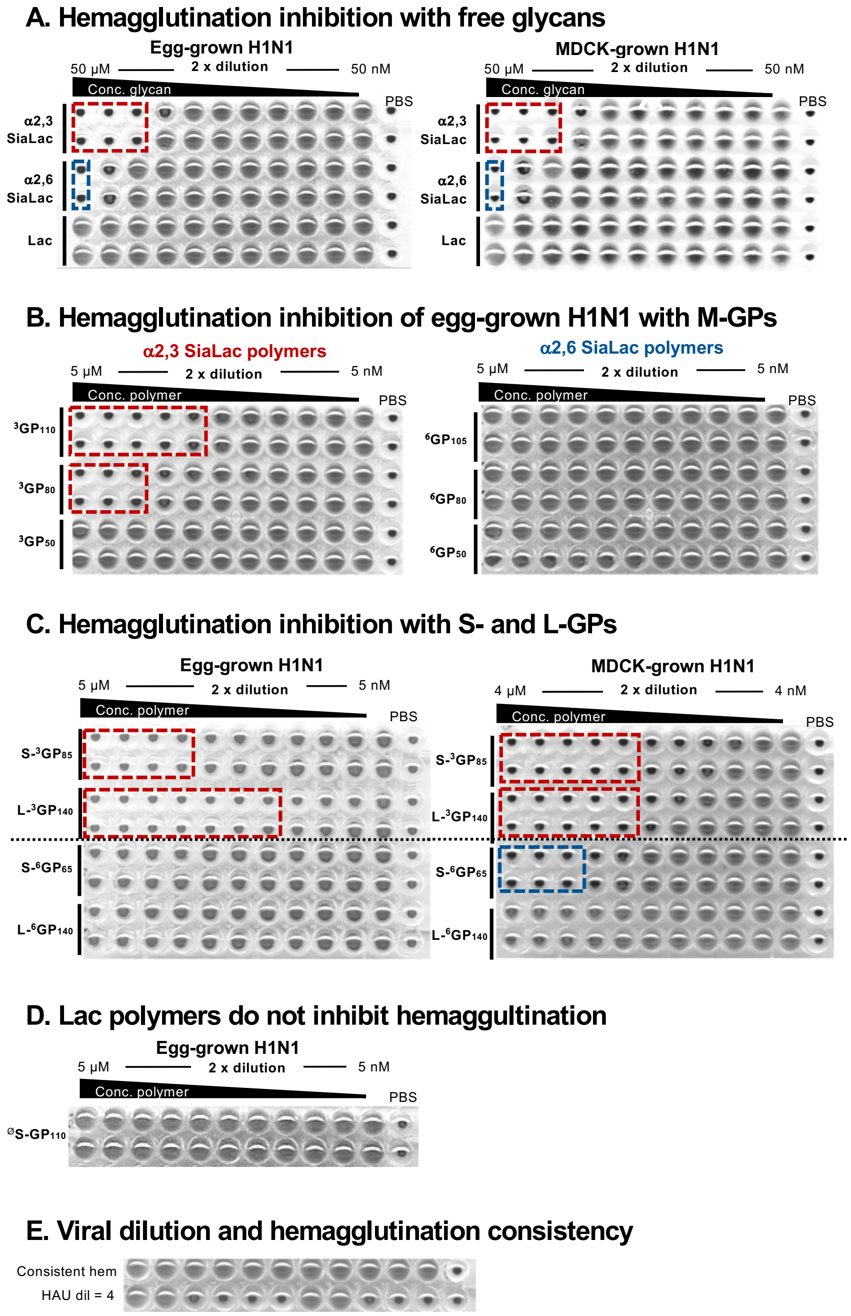

**Figure S10. Hemagglutination inhibition assays.** (A) The free sialyllactose glycans inhibit egg-grown and MDCK-grown H1N1 to the same extent. (B) The valency dependence of M-$^{3}$GPs can be seen for the egg-grown H1N1 ($K_i$ of M-$^{3}GP_{110}$ = 313 nM, $K_i$ of M-$^{3}GP_{80}$ = 1.25 µM, and $K_i$ of M-$^{3}GP_{50}$ > 5 µM) while the M-$^{6}$GPs do not inhibit hemagglutination at any valency. (C) Short and long $^{3}$GPs inhibit hemagglutination of the egg grown virus $K_i$ of S-$^{3}GP_{85}$ = 625 nM and $K_i$ of L-$^{3}GP_{140}$ = 78 nM. The MDCK virus is inhibited to equal extent by the short and long $^{3}$GPs, $K_i$ = 234 nM and is also inhibited by S-$^{6}GP_{65}$, $K_i$ = 938 nM. (D) Lactose polymers do not inhibit hemagglutination, shown for the egg-grown H1N1 with $^{\varnothing}$S-$GP_{110}$. (E) For all hemagglutination assays, a viral dilution with HAU = 4 was used and shown to be consistent across replicates.

**Table S5. Hemagglutination inhibition constants.** The HI constants are shown for each virus type and inhibitor used. Where inhibition was not seen, the value in the table was entered as the highest concentration used.

| Virus | Inhibitor | $K_i$ (μM) |
|---|---|---|
| H1N1 EGG | **α2,3-SiaLac** | 13.0 |
| H1N1 EGG | **α2,6-SiaLac** | 50.0 |
| H1N1 EGG | **Lac** | > 50.0 |
| H1N1 MDCK | **α2,3-SiaLac** | 13.0 |
| H1N1 MDCK | **α2,6-SiaLac** | 50.0 |
| H1N1 MDCK | **Lac** | > 50.0 |
| H1N1 EGG | **M-$^3$GP$_{110}$** | 0.3 |
| H1N1 EGG | **M-$^3$GP$_{80}$** | 1.3 |
| H1N1 EGG | **M-$^3$GP$_{50}$** | >5.0 |
| H1N1 EGG | **M-$^6$GP$_{105}$** | >5.0 |
| H1N1 EGG | **M-$^6$GP$_{80}$** | >5.0 |
| H1N1 EGG | **M-$^6$GP$_{50}$** | >5.0 |
| H1N1 EGG | **S-$^3$GP$_{85}$** | 0.6 |
| H1N1 EGG | **L-$^3$GP$_{140}$** | 0.08 |
| H1N1 EGG | **S-$^6$GP$_{65}$** | >5.0 |
| H1N1 EGG | **L-$^6$GP$_{140}$** | >5.0 |
| H1N1 MDCK | **S-$^3$GP$_{85}$** | 0.2 |
| H1N1 MDCK | **L-$^3$GP$_{140}$** | 0.2 |
| H1N1 MDCK | **S-$^6$GP$_{65}$** | 0.9 |
| H1N1 MDCK | **L-$^6$GP$_{140}$** | >4.0 |

### H1N1 and H3N2 binding to density variant mucin-mimetic arrays

H1N1 and H3N2 viruses were isolated and utilized neat from embryonated chicken eggs allantoic fluid or MDCK viral culture supernatant by incubating on the array for 1 hour at room temperature with rocking. The slide was washed two times with BSA/PBST. H1N1 was fixed to the array with a 20 minute incubation with 4% PFA. This was followed by an incubation of a 1:500 dilution of anti-HA in BSA/PBST on the array for 1 hour at room temperature with rocking. For H3N2, the virus was fixed on the array with a solution of 4% PFA for 10 minutes. Then the membrane was permeabilized in 70% ethanol for 10 min so that a 1:500 dilution of NP antibody could be incubated on the array for 1 hour at room temperature with rocking. Following primary antibody incubation, the slides were washed two more times with BSA/PBST and a 1:500 dilution of anti rabbit-AF647 antibody was incubated in the dark with shaking. The subarrays were washed two more times with BSA/PBST, two times with the 0.1% Tween-20 solution, rinsed with MilliQ, spin dried, and imaged at the highest PMT possible without producing saturated pixels.

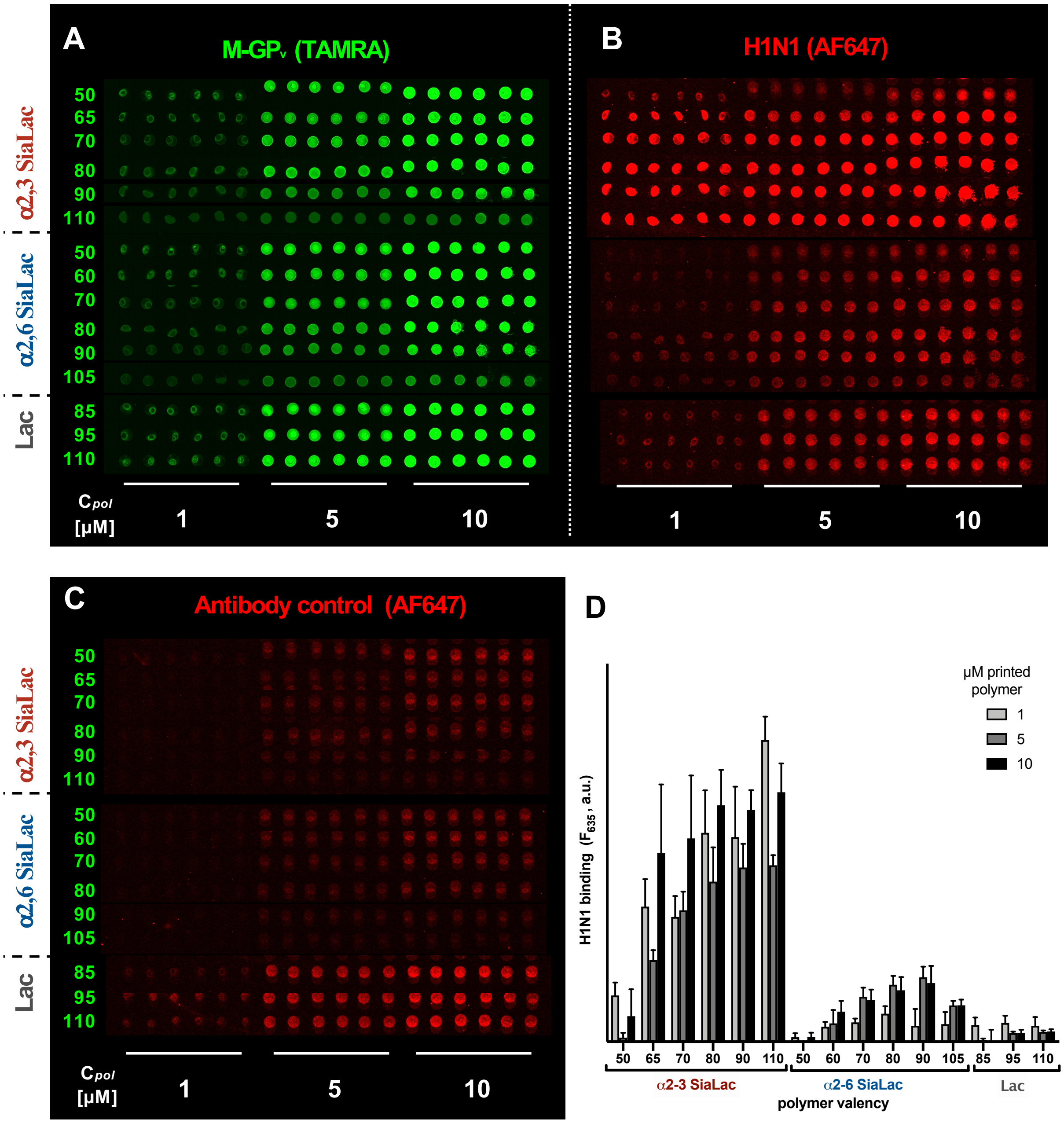

**Figure S11. Full array fluorescence scan and bar graph representation of egg H1N1 binding in mucin-mimetic arrays containing M-GPvs (extended data for Figures 4A and B).**
(A) Fluorescence scan ($\lambda_{em}$=532 nm) of density variant array comprising TAMRA-labeled medium (**M**) glycopolymers carrying α2,3-sialyllactose (**$^{3}$GP**), α2,6-sialyllactose (**$^{6}$GP**), or lactose (**$^{Ø}$GP**) printed at 1, 5 or 10 µM concentrations. (B) Fluorescence scan ($\lambda_{em}$ = 635 nm) of mucin mimetic array incubated with H1N1 and stained with AF647-labeled anti-HA antibody. (C) Fluorescence scan ($\lambda_{em}$ = 635 nm) of anti-HA antibody background binding. (D) Bar graph representation of H1N1 binding intensity in mucin mimetic arrays after background signal (anti-HA only) subtraction. Values represent mean values for six replicate spots. Error bars represent standard deviation from 6 replicate spots.

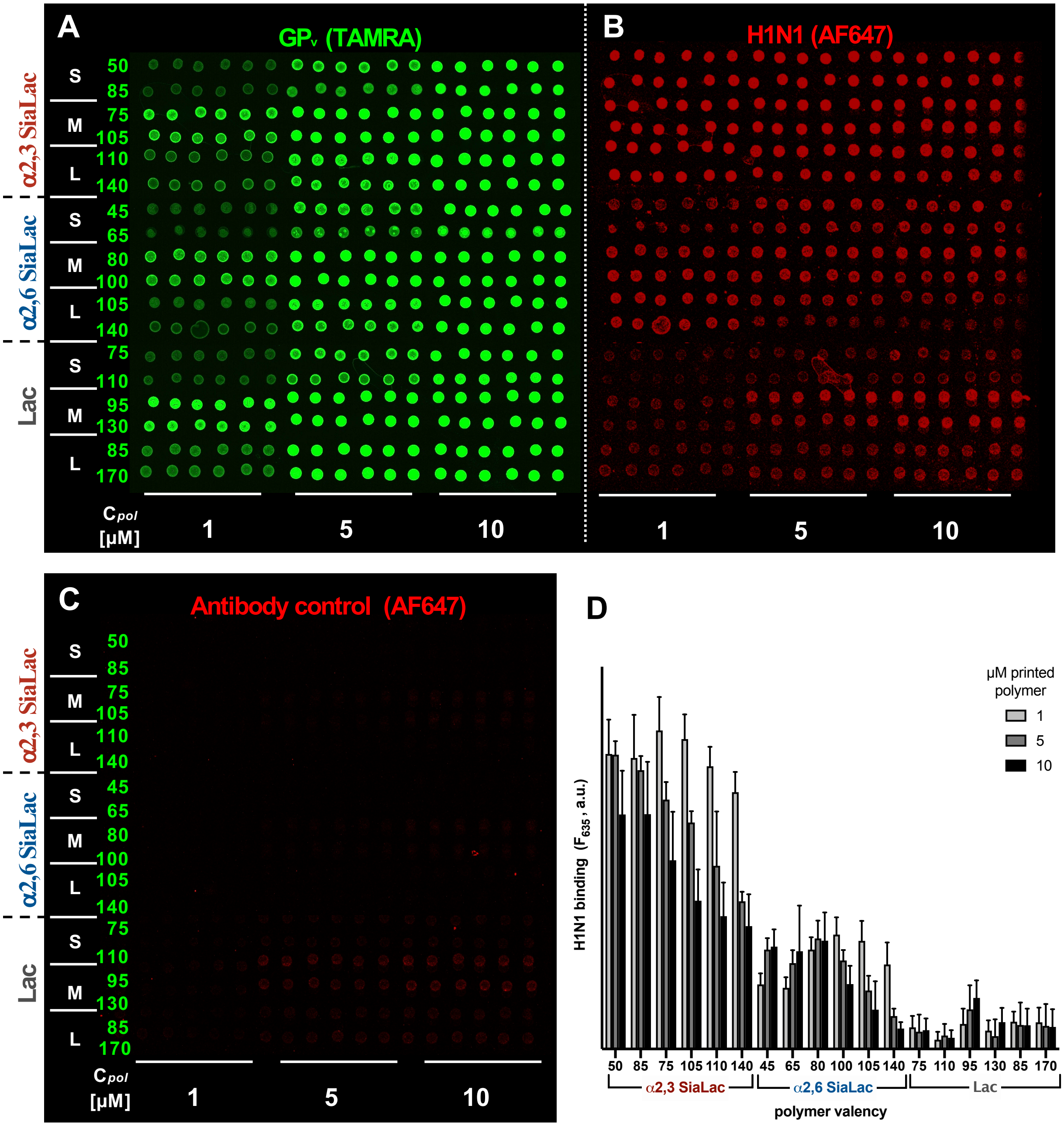

**Figure S12. Full array fluorescence scan and bar graph representation of egg H1N1 binding in mucin-mimetic arrays of all polymer lengths (extended data for Figures 4C and D).**
(A) Full microarray print containing **S/M/L-GPs** at half and full glycosylation printed in six replicates at 1, 5, and 10 μM. (B) H1N1 from egg bound to the array and probed with anti-HA primary antibody and AF647 labeled secondary antibody. (C) The antibody control contains both the HA primary antibody and labeled secondary and shows some background binding to the Lac polymers. (D) The signal from the antibody control subarray is subtracted from the H1N1 array before plotting viral binding.

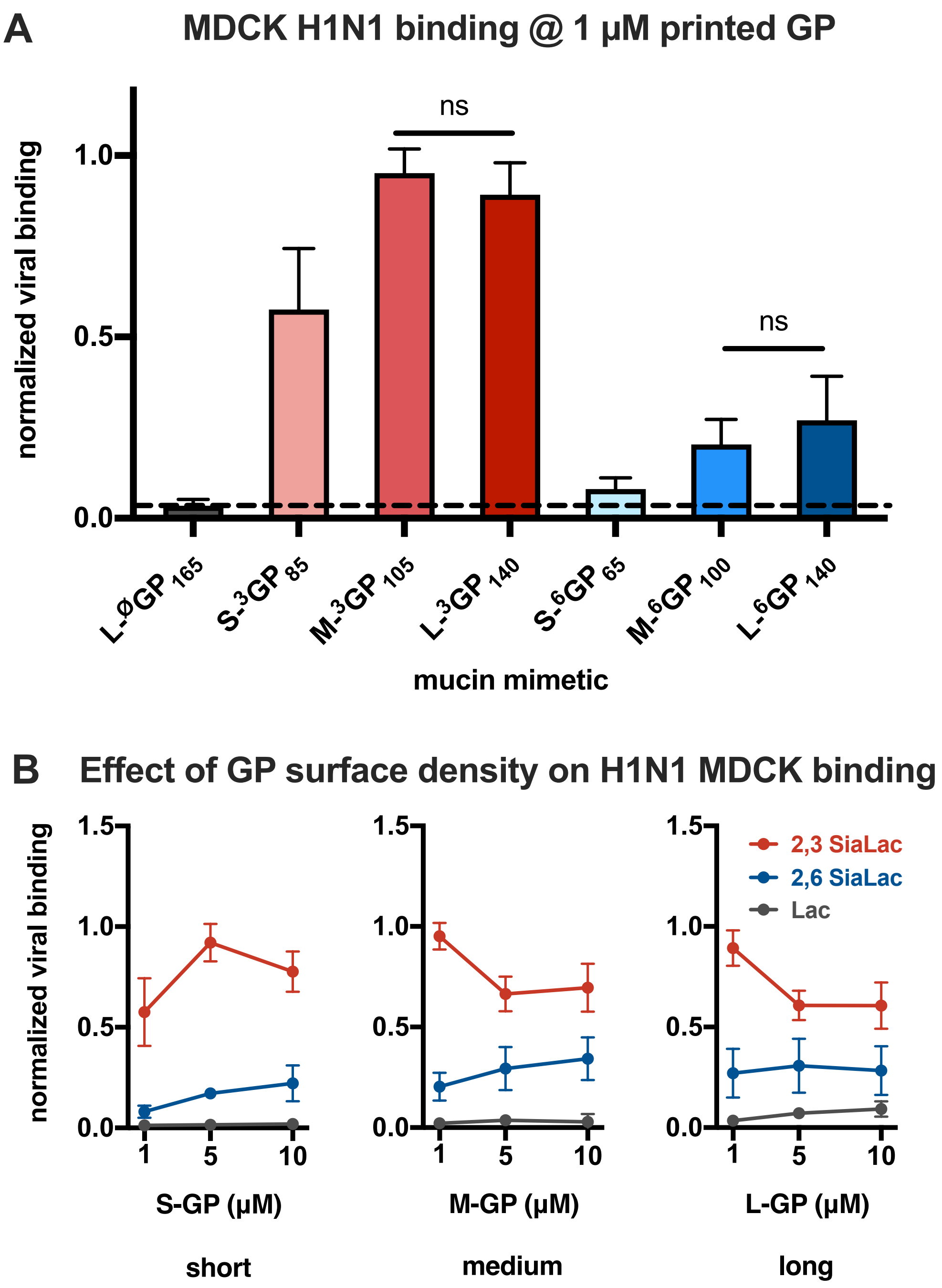

**Figure S13. MDCK produced H1N1 binding to mucin mimetic arrays.** (A) Normalized binding of H1N1-MDCK to all glycopolymer lengths a 1 μM printing concentration. The dotted line signifies the Lac background. (B) The effect of **GP** printing concentration on MDCK produced H1N1 binding

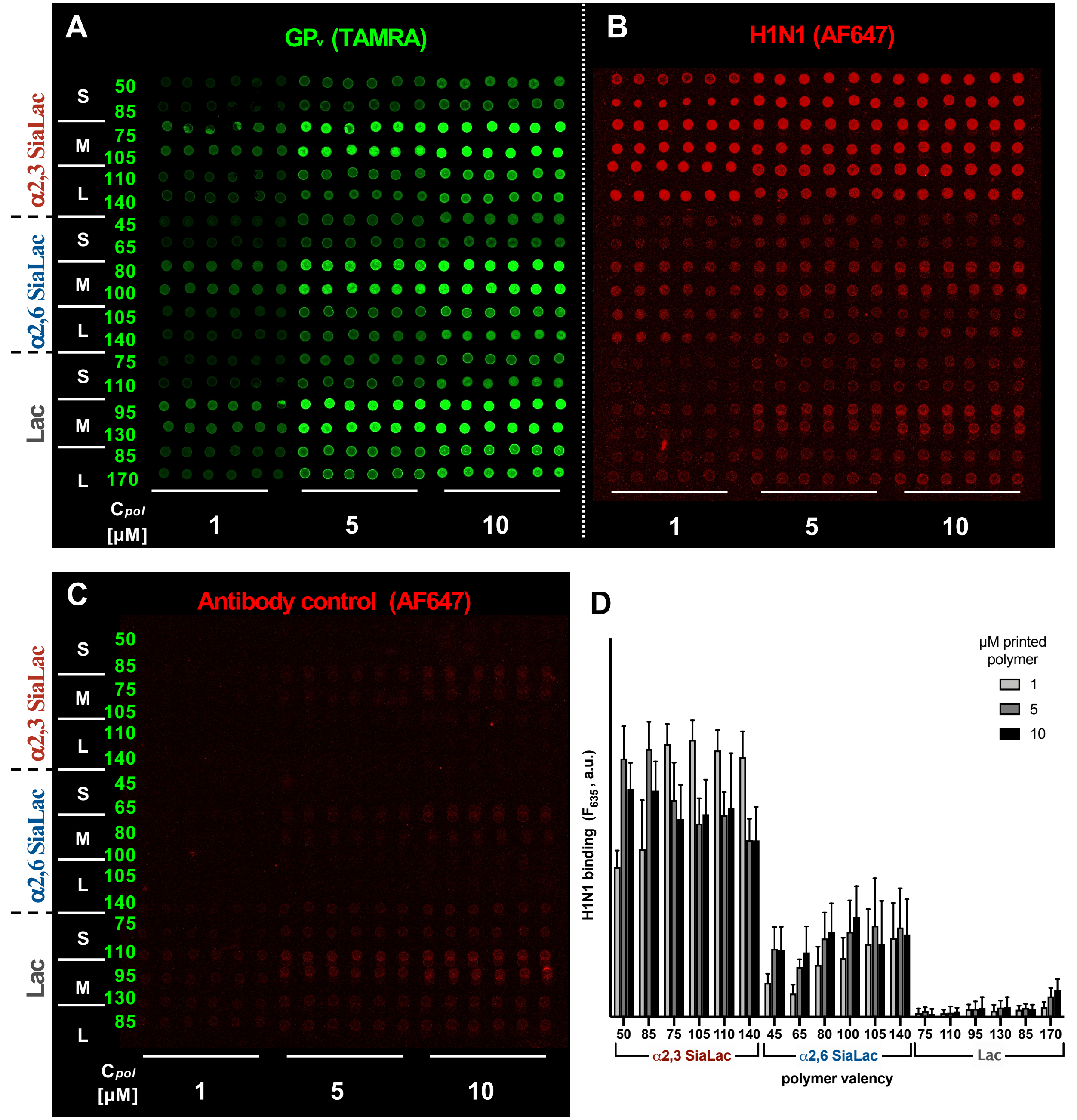

**Figure S14. Full array fluorescence scan and bar graph representation of H1N1 MDCK binding in mucin-mimetic arrays of all polymer lengths.** (A) Full microarray print containing **S/M/L-GPs** at half and full glycosylation printed in six replicates at 1, 5, and 10 µM. (B) H1N1 from MDCK cells bound to the array and probed with anti-HA primary antibody and AF647 labeled secondary antibody. (C) The antibody control contains both the HA primary antibody and labeled secondary and shows some background binding to the Lac polymers. (D) The signal from the antibody control subarray is subtracted from the H1N1 array before plotting viral binding.

### SVM analysis

#### Machine learning analysis of H1N1 array binding data

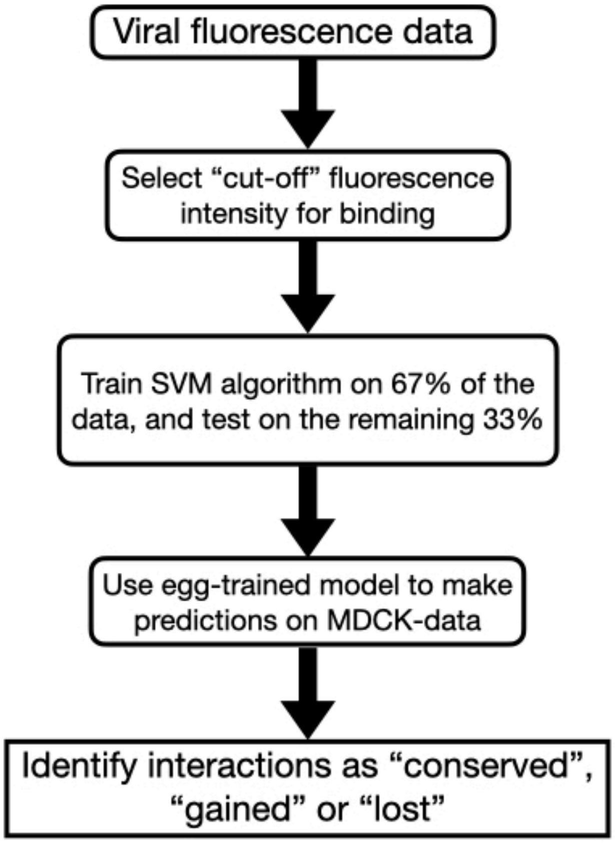

**Figure S15. Machine learning workflow used in this work.** (A) Data preparation for each (H1N1 EGG and H1N1 MDCK) dataset, as described in Materials and Methods. Viral binding fluorescence data for each data set was used to train separate SVM algorithms. (B) Model trained on H1N1 EGG data was then used to make predictions on the H1N1 MDCK data (“cross-model” prediction), and the results were compared with those from the model trained on H1N1 MDCK (“self-model” prediction). Based on the similarities and differences in the outcomes of the two models, data points were categorized as “Conserved”, “Gained in MDCK”, and “Lost in MDCK”, as described in Materials and Methods and Figure S19.

**Establishing fluorescence threshold for binding**

Since the virus is known to be a non-binder to lactose, the lactose fluorescence was used as negative control. Based on the distribution of normalized fluorescence intensities of all 3 types of glycans (lactose, 2,3-SiaLac and 2,6-SiaLac), the cutoff was determined to be the least normalized fluorescence value for which lactose fluorescence was negligible.

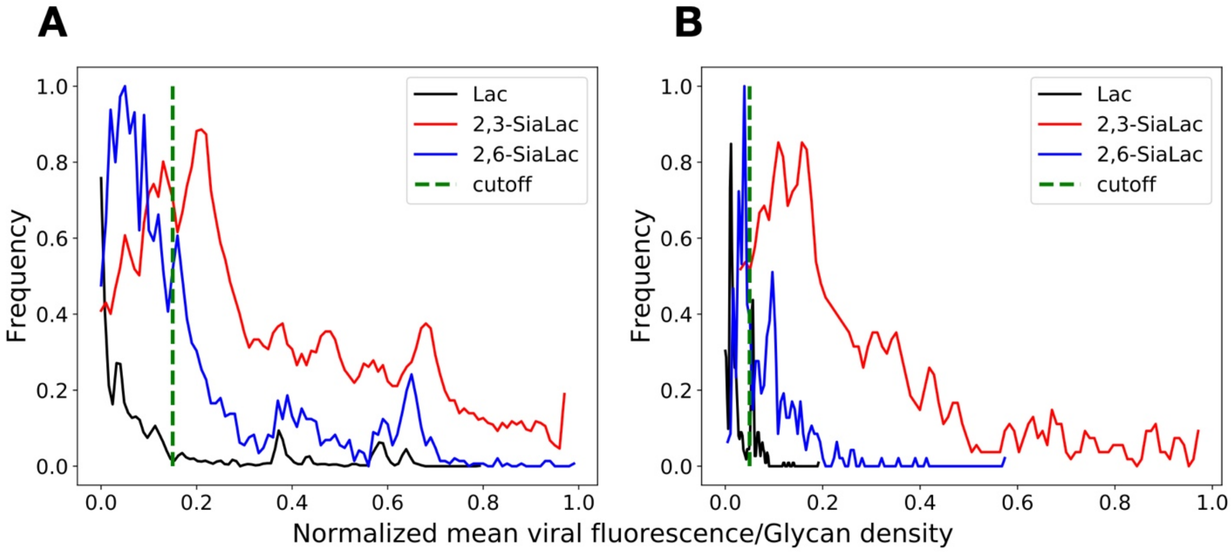

**Figure S16. Establishing fluorescence threshold for binding.** Distribution of fluorescence for lactose (black), 2,3-SiaLac (red), and 2,6-SiaLac (blue) are shown for (A) H1N1 EGG and (B) H1N1 MDCK data sets. Cutoff value (broken green line) was determined to be 0.15 fo H1N1 EGG and 0.05 for H1N1 MDCK.

**Convergence of SVM training**

Performance of binary classification, namely, accuracy, precision, recall, and F-1 score, were defined as:

$$accuracy = \frac{True\ positive + True\ negative}{All\ data}$$

(fraction of correctly classified data, calculated using the following equation)

$$precision = \frac{True\ positive}{True\ positive + False\ positive}$$

(fraction of total predicted positives that are true)

$$recall = \frac{True\ positivee}{True\ positive + False\ negative}$$

(fraction of all positives that are correctly predicted)

$$F1\ score = 2 * \frac{Precision * Recall}{Precision + Recall}$$

(balance between precision and recall)

Convergence of SVM training was defined when these quantities no longer changed with further iterations.

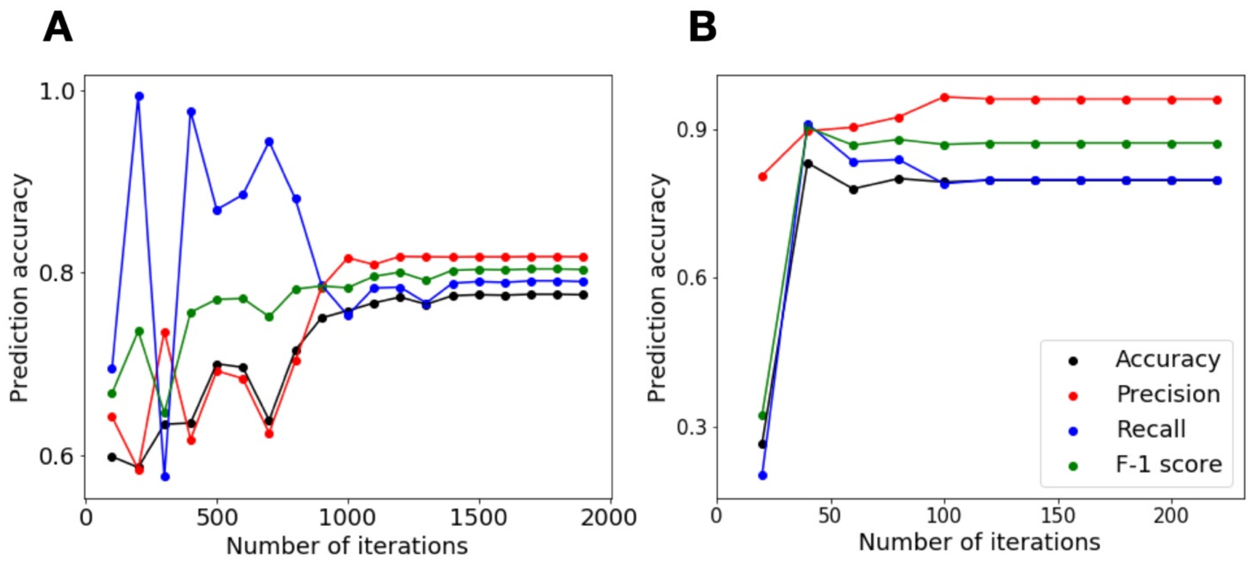

**Figure S17. Convergence of SVM training.** Prediction accuracy (black), Precision (Red), Recall (Blue) and F-1 score (Green) are shown as a function of number of iterations, for (A) H1N1 EGG and (B) H1N1 MDCK data sets. Training with the former data converged in 2000 iterations, while that of the latter converged in 200 iterations.

**Validation of SVM training**

SVM algorithim trained on the training data set (66% randomly selected data points from the data set) was used to predict the remaining, testing data set (33% of the data set). The results are shown in the form of a confusion matrix where the X-axis is the prediction output, and the Y-axis is the experimental outcome. Number of correct predictions are shown across the diagonal (correctly predicted non-binding shown on top left, correctly predicted binding shown on bottom right). False positives (predicted binding for points that showed non-binding in experiment) are shown top right. False negatives (predicted non-binding for points that showed binding in experiment) are shown on bottom left.

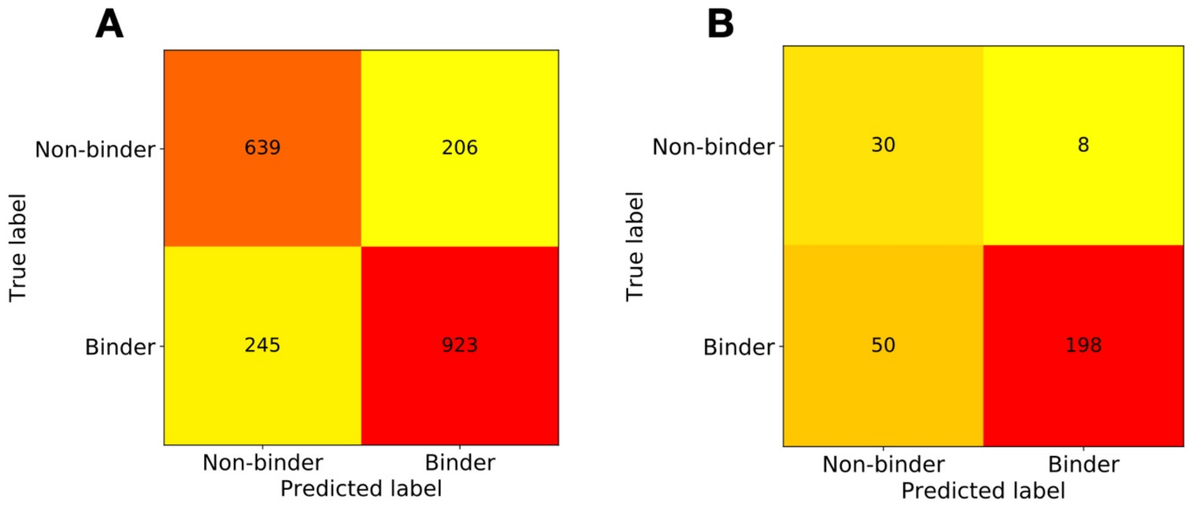

**Figure S18. Validation of SVM testing.** Confusion matrix for the prediction of testing data for (A) H1N1 EGG and (B) H1N1 MDCK data sets.

**Comparison of interaction patterns**

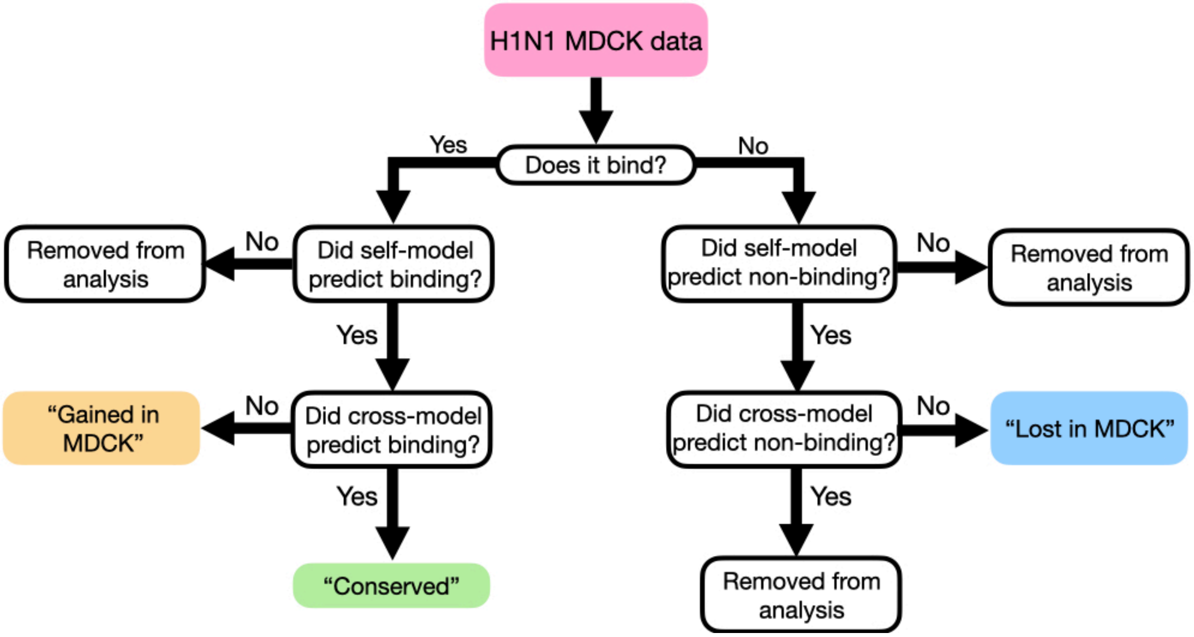

**Figure S19. Comparison of interaction patterns.** True binders (left branch) that were correctly predicted by both self- and cross-models were termed "Conserved". Those that were correctly predicted by the self-model but not by the cross-model (implying that these interactions did not exist in H1N1 EGG) were termed "Gained in MDCK". Those that were wrongly predicted by the self-model were not used in this analysis. True non-binders (right branch) that were correctly predicted by the self-model but not by the cross-model (implying that these interactions existed in H1N1 EGG but are absent in H1N1 MDCK) were termed "Lost in MDCK".

### Western blot analysis of PNGase treated of H1N1

To prepare samples for PNGase treatment, 6 µL of IAV were combined with 1 µL of Glycoprotein Denaturing Buffer and 3 µL of MilliQ in PCR tubes. Samples were denatured at 100 $^{o}$C for 10 min, then chilled on ice and centrifuged briefly (10 sec). The reaction volume of each sample was adjusted to 20 µL by adding GlycoBuffer 2 (2 µL), 10% NP-40 (2 µL), and of MilliQ (6 µL). Then 1 µL of PNGase (500 units) was added and the tubes were mixed gently (1 µL of MilliQ was added to non-PNGase treated samples). The samples were incubated at 37 $^{o}$C for 4 hr and placed at 4 $^{o}$C until running the gel.

To prepare the samples for the gel, an equal volume (20 µL) of 2.5% loading dye with 0.4 M DTT was added and they were incubated at 95 $^{o}$C for 10 min. A 4-12% Bis Tris gel was loaded with 15 µL of sample (8 µL of ladder was loaded to terminal wells). The gel was run at a constant 200 V for 22 min in 1x MES buffer, after which time it was rinsed with water and transferred to a PVDF membrane using the P3 settings (20 V for 7 min) on the iBlot2.

Following transfer, the membrane was block with a 5% blotto solution in PBST (0.1% Tween-20) for 1 hr at room temperature. Following rinses with PBST, the membrane was probed overnight at 4 $^{o}$C with a 1:500 dilution of anti-HA antibody in PBST. The following day, it washed for 15 min (3 x 5 min) with PBST and incubated with a 1:2000 dilution of anti-rabbit IgG, HRP-linked antibody for 1 hr at room temperature. The membrane was washed again for 15 min (3 x 5 min) in PBST. The HRP substrate was added for 30 sec and the blot was imaged on a BioRad GelDoc XRS+ imager.

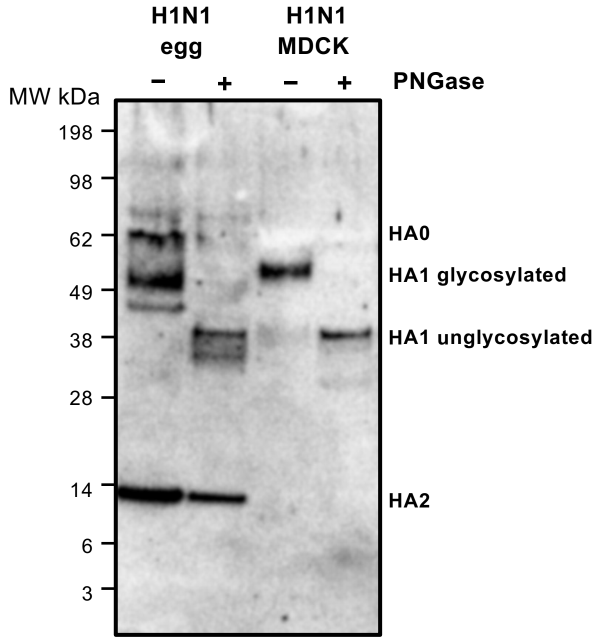

**Figure S20. Western blot analysis of *N*-glycosylation of HA from H1N1.**
The HA of the egg grown virus is slightly smaller than the HA from the MDCK grown virus. Removal of N-glycans through PNGase treatment yields HAs of the same molecular weight.
